# The trade-off between individual metabolic specialization and versatility determines the metabolic efficiency of microbial communities

**DOI:** 10.1101/2023.04.28.538737

**Authors:** Miaoxiao Wang, Xiaoli Chen, Yuan Fang, Xin Zheng, Ting Huang, Yong Nie, Xiao-Lei Wu

## Abstract

In microbial systems, a metabolic pathway can be either completed by one autonomous population or alternatively be distributed among a consortium performing metabolic division of labor (MDOL), where several specialized populations cooperate to complete the pathway. MDOL facilitates the system’s function by reducing the metabolic burden; however, it may also hinder the function by reducing the exchange efficiency of metabolic intermediates among individuals. As a result, the metabolic efficiency of a community is influenced by trade-offs between the metabolic specialization and versatility of individuals, with the latter potentially introducing metabolic redundancy into the community. However, it remains unclear how metabolic specialization and versatility of the individuals involved can be controlled in order to optimize the function of the community. In this study, we deconstructed the metabolic pathway of naphthalene degradation into four specialized steps and introduced them individually or combinatorically into different strains, with varying levels of metabolic specialization. Using these strains, we engineered 1,456 synthetic consortia with varying levels of metabolic redundancy and tested their naphthalene degradation efficiency. We found that 74 consortia possessing metabolic redundancy exhibited higher degradation efficiency than both the autonomous population and the rigorous MDOL community. Quantitative modeling derived from our experiments provides general strategies for identifying the most effective MDOL consortium with functional redundancy (MCFR) from a range of possible MCFRs. Our large-scale genomic analysis suggests that natural communities for hydrocarbon degradation are mostly functionally redundant. In summary, our study provides critical insights into the engineering of high-performance microbial systems and explains why functional redundancy is prevalent in natural microbial communities.

## Introduction

Microorganisms colonize all major ecological niches on our planet, where they accomplish complex metabolic tasks that are critical for survival in often harsh environments^1,2^. These tasks, including carbon fixation, nitrogen cycling, sulfur oxidation/reduction, methanogenesis, and breaking down organic matter, are critical for maintaining the delicate balance of global biogeochemical cycles and play a fundamental role in sustaining our planet’s ecosystems^1,3^. In nature, these metabolic processes are accomplished through long metabolic pathways. It is therefore of paramount importance to disseminate how natural microorganisms develop strategies that result in efficient pathways. Such understanding should greatly advance rational engineering as well as the *de novo* construction of metabolic pathways, thus benefiting diverse industries, including biomanufacturing^4,5^, biomedicine^6,7^, and bioremediation^8,9^.

Microorganisms reside and interact with myriads of other species, forming complex microbial communities. Within a community, a metabolic pathway can be completed by one single population that can autonomously produce all the enzymes required for a pathway (Figure 1A). Alternatively, the pathway can be cooperatively completed by several interacting populations, with each of its microbial strains performing one specific metabolic step of the pathway (i.e., it is metabolically specialized), a phenomenon termed metabolic division of labor^10-13^ (MDOL; we termed such systems rigorous MDOL; Figure 1B). In terms of achieving high pathway productivity, both autonomous population and MDOL systems are characterized by both advantages and disadvantages (Figure 1C). First, while all individuals residing within an autonomous population are able to perform the entire metabolic pathway, only a subset of individuals in a rigorous MDOL system can perform specific steps. Therefore, the average functional capacity of the rigorous MDOL system to perform each step is potentially lower than that in the system containing autonomous populations. Assuming that both systems maintain the same population size, the system containing autonomous populations may possess a higher functional capacity in each step of the whole pathway. Second, if every enzyme involved in the complete metabolic pathway is expressed in a single strain, it might result in a significant metabolic burden on that single population, potentially leading to a reduction in total biomass and overall productivity^14^. In comparison, each population in an MDOL system only produces a subset of enzymes required for the overall pathway, thereby diminishing the metabolic burden experienced by each population^10,12,13^. Third, the intermediate metabolites are exchanged across different populations in an MDOL system. In many microbial systems, such exchange is mediated by passive diffusion with an unavoidable metabolite loss and lower exchange efficiency^15-17^. As a result, the inefficiency of intermediate exchange is likely to limit the function of an MDOL system.

**Figure 1.**
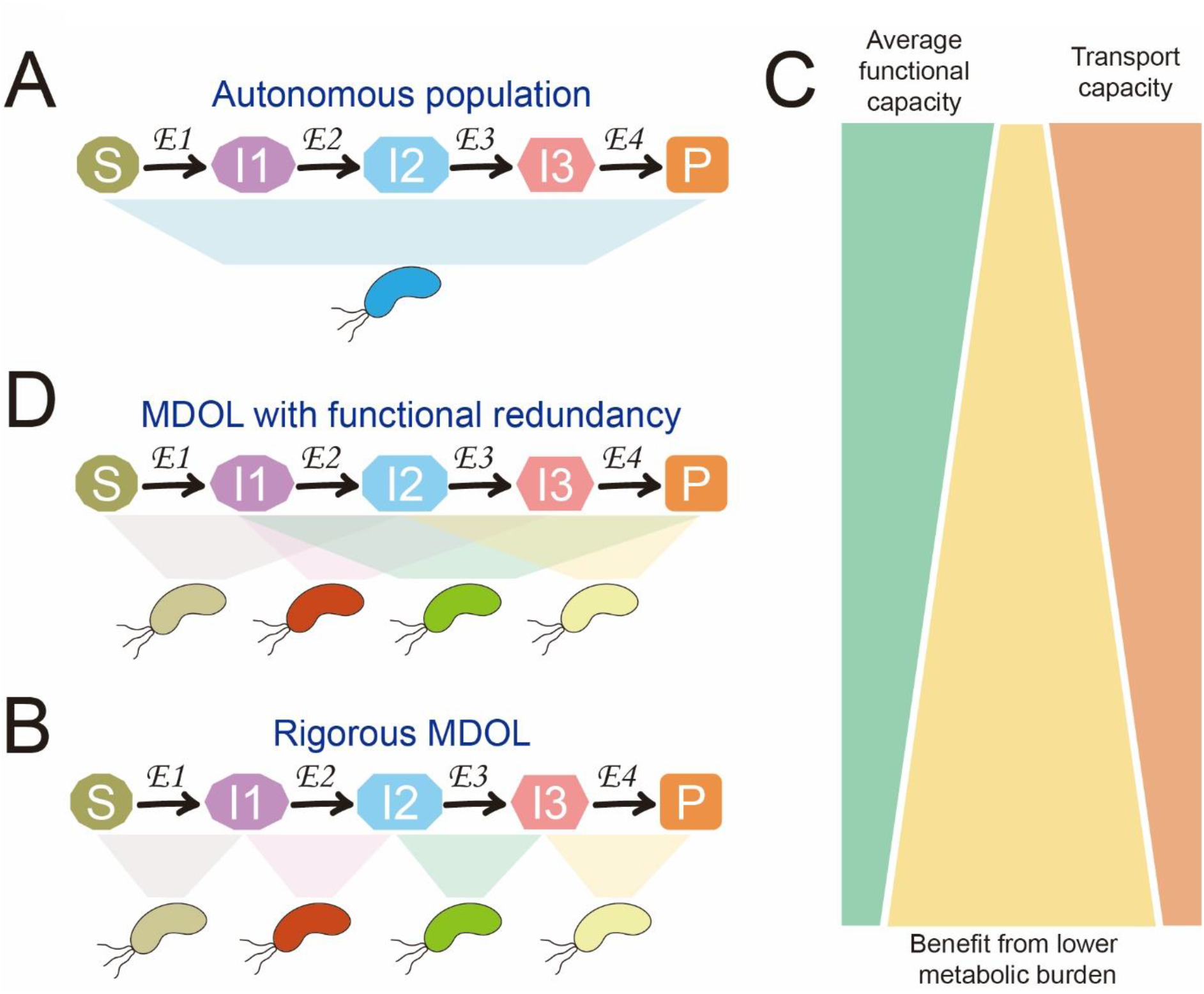
Schematic diagram of different microbial systems performing a metabolic pathway. (A) An autonomous population: one single population that can produce all the enzymes required for a pathway so it can autonomously execute the whole pathway. (B) A rigorous metabolic division of labor (MDOL) consortium: a consortium composed of several interacting populations to collectively execute the pathway, each of which can only perform a specific metabolic step of the pathway. (D) An MDOL consortium possessing functional redundancy (MCFR): a consortium composed of several interacting populations to collectively execute the pathway, each of which can only perform more than one metabolic step so each step in the pathway can be redundantly performed by multiple populations. Note that the authors only provide a typical example of MCFR. More configurations are summarized in Figure S1. (C) The hypothesized features of the proposed three systems, including average functional capacity and the average metabolic burden of each individual, as well as the overall efficiency of intermediate transport. Among the three types of microbial systems, the autonomous population possesses the highest average functional capacity and the overall efficiency of intermediate transport, which benefit its function. It also possesses the highest average metabolic burden that harms its function. The rigorous MDOL consortium possesses opposite traits against the autonomous population. MCFRs possess the intermediate level of these traits between the autonomous population and the rigorous MDOL system.

If one can only choose either an autonomous population or a two-step MDOL system (i.e., when a pathway can only segregate into two steps) for metabolic pathway optimization, a trade-off decision can be made based on the metabolic burden and exchange efficiency, as reported by a recent study^13^. When a pathway can segregate into more than two steps, it should be possible to adopt a third strategy by introducing functional redundancy to a rigorous MDOL system (Figure 1D). Functional redundancy is defined as the coexistence of multiple distinct taxa capable of performing the same focal biochemical function^18,19^. An MDOL consortium possessing functional redundancy (MCFR) denotes that one population in a consortium can perform more than one metabolic step (i.e., it is metabolically versatile) and each step in the pathway may be redundantly executed by multiple populations. Because MCFRs represent the intermediate states between the autonomous population and the rigorous MDOL system (Figure 1C), we propose that MCFRs combine the advantages of both systems. Specifically, MCFRs comprised of versatile populations can maintain functional redundancy of the metabolic steps, thus ensuring the high average functional capacity of the community and rendering the exchange of metabolic intermediates highly efficient. In addition, by avoiding the production of all enzymes in one population, MCFRs can reduce the metabolic burden on individual populations. Therefore, we hypothesize that MCFRs show higher function than both systems of the autonomous population and the rigorous MDOL.

In this study, we set out to test our hypothesis by engineering synthetic consortia with different functional redundancy for naphthalene degradation. Based on our experimental results, we built a mathematical model to explore the general strategies used for constructing a high-function community from the individuals with appropriate levels of metabolic specialization.

## Results

### Synthetic consortia with functional redundancy exhibit a higher function of naphthalene degradation

To test our hypothesis, we first built synthetic consortia that perform metabolic division of labor and possess different levels of functional redundancy (MCFRs) and compared the function of these consortia to the corresponding autonomous population and the rigorous metabolic division of labor (MDOL) system. To this end, we engineered an autonomous *Pseudomonas stutzeri* strain that completely degrades naphthalene via a linear pathway (Methods; Supplementary Information S1.1; Figure S1). Next, we partitioned the naphthalene degradation pathway into four steps and engineered fourteen strains that perform either one specific or multiple metabolic step(s). We conceptualized the genotypes of these strains using bit strings containing “0” and “1”. A “0” in a certain bit position of the string indicates that the corresponding genotype lacks the gene responsible for the specific step at that position; a “1” denotes that one genotype contains genes encoding the enzymes for a step (Figure S1 and Figure S2A). For example, strain [0, 1, 1, 0] indicates that it can perform the second and third steps but not the first and fourth steps. As summarized in Figure S2B, co-culturing two, three, or four of these fourteen strains would generate either 91, 364, or 1001 possible synthetic consortia for naphthalene degradation, respectively. Of those consortia, we found that respectively 25, 230, or 861 consortia possess the complete degradation pathway (i.e., one metabolic step can be at least performed by one strain in the consortium). Furthermore, respectively 18, 224, or 860 of these consortia possessed functional redundancy (i.e., at least one metabolic step is performed by more than one strain in the consortium). To quantify the function of these consortia, we measured the naphthalene degradation rates and growth rates of all the possible two-, three- and four-member consortia.

When we assessed the naphthalene degradation rates of the two-member MCFRs, we found that four consortia, that is, the consortia composed of [1, 1, 1, 0] & [0, 0, 1, 1], [1, 1, 1, 0] & [0, 1, 0, 1], [1, 1, 1, 0] & [1, 0, 0, 1], and [1, 1, 0, 1] & [0, 1, 1, 1], exhibited higher degradation rates than the two-member rigorous MDOL consortia (i.e., [1, 1, 1, 0] & [0, 0, 0, 1], [1, 1, 0, 0] & [0, 0, 1, 1], [1, 0, 0, 0] & [0, 1, 1, 1]; Figure S3A). In particular, we found that the function of the consortium [1, 1, 1, 0] & [0, 1, 0, 1] was even better than that of the autonomous population [1, 1, 1, 1] (53.8% ± 0.3% versus 45.3 ± 0.5%, *p* < 0.001). When we included more members in our MCFRs, we identified many consortia that exhibited degradation rates than both the autonomous population and the corresponding rigorous MDOL system, including ten three-member MCFRs (Figure S3B), and 63 four-member MCFRs (Figure 2A-B). In particular, seven four-member MCFRs possessed a degradation rate of over 80 % within four days (Figure 2B), which was significantly higher than the autonomous population (showing a degradation rate of 45.3 ± 0.5%), and the four-member rigorous MDOL system ([1, 0, 0, 0] & [0, 1, 0, 0] & [0, 0, 1, 0] & [0, 0, 0, 1] with a degradation rate of only 10.6 ± 2.2 %). Among these MCFRs, the consortium composed of [1, 0, 1, 1] & [1, 1, 0, 0] & [1, 1, 0, 1] & [1, 1, 1, 0], exhibit the highest degradation rate (83.8 ± 0.2%). We obtained similar results when we compared the overall growth rates of these consortia (Figure S3C-D; Figure S4A). We found that 12 three-member MCFRs (Figure S3D), and 54 four-member MCFRs (Figure S4B) grew faster than both the autonomous population and the rigorous MDOL system. Furthermore, we found that the growth rates of the consortia directly correlated with their degradation rates (Figure S5). Together, these results clearly suggested that MDOL systems characterized by appropriate levels of functional redundancy are better suited to efficiently complete entire pathways.

**Figure 2.**
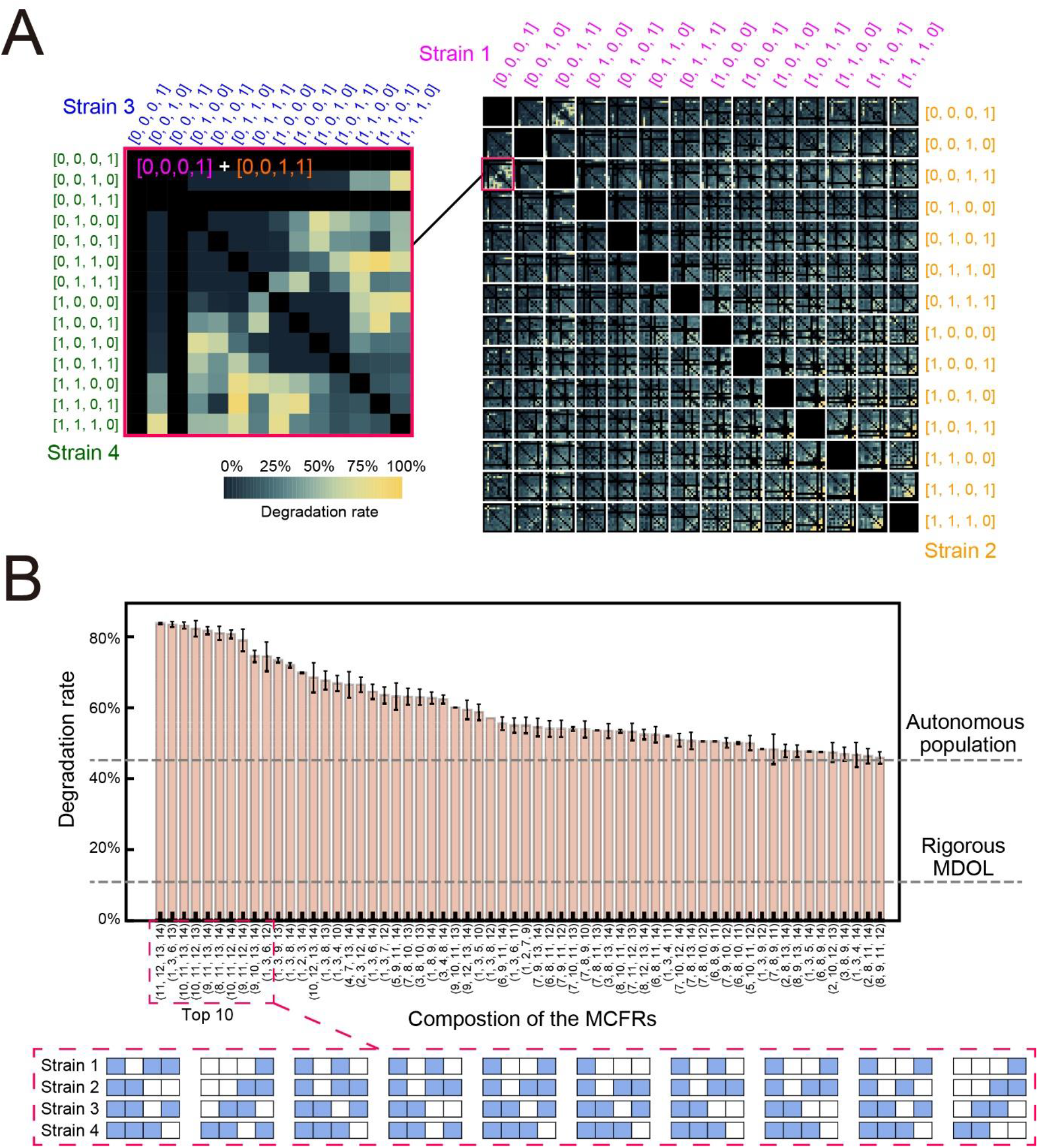
Degradation rates of the four-member consortia. (A) 14×14×14×14 matrix diagrams show the naphthalene degradation rates of the four-member consortia. Each 14×14 panel correspond to a fixed strain 1 (violet) and a fixed strain 2 (orange) against all combination of strains 3 (blue) and 4 (green). The color intensity represents the naphthalene degradation rate after 96-h culture, and the value shown is the average value of the three experimental replicates. (B) Summary of the high-function consortia. Naphthalene degradation rates of 63 high-function consortia are shown, which are significantly higher than those of the autonomous population and the rigorous metabolic division of labor (MDOL) consortium (Indicated by the grey dashing lines; Double-tailed Student’s T-test, *p* <0.05). Data were calculated from three independent replicates. Each consortium is named after a four-number vector, in which each number represents the decimal form of the bit string of one member genotype of the consortium, as follows: 1 – [0, 0, 0, 1], 2 – [0, 0, 1, 0], 3 – [0, 0, 1, 1], 4 – [0, 1, 0, 0], 5 – [0, 1, 0, 1], 6 –[0, 1, 1, 0], 7 – [0, 1, 1, 1], 8 – [1, 0, 0, 0], 9 – [1, 0, 0, 1], 10 – [1, 0, 1, 0], 11 – [1, 0, 1, 1], 12 – [1, 1, 0, 0], 13 – [1, 1, 0, 1], 14 – [1, 1, 1, 0]. In the bottom graph, the composition of the consortia possessing the top ten performances is visualized. The composition of each consortium is indicated by a four-row array. Each row represents the genotype of one strain involved in the consortium to perform the four-step, where the blue grid indicates that the strain possesses can perform the corresponding step while the white grid indicates the strain is unable to perform that step.

### The functions of MCFRs can be predicted by the metabolic burden of strains, average functional capacity, and transport capacity of metabolites

We hypothesized that the MCFRs exhibit higher function because their configurations better mediate the conflicts among the metabolic burden of strains, average functional capacity per biomass unit, as well as their capacity to transport metabolites. If this explanation is correct, then the functions of MCFRs, autonomous population, and rigorous MDOL should be largely determined by these three factors. We tested this duction by using the measured degradation rates and growth rates of our four-member MCFRs, as well as by defining four quantitative indices for each consortium: (1) functional redundancy level (FR) is defined based on functional dissimilarities among species^19^ (Supplementary S1.3); (2) the average metabolic burden (AMB) represents the average value of metabolic burdens of the strains involved in the consortium; (3) The average functional capacity (AFC) represents the average value of the functional capacity of the strains involved in the consortium; (4) The transport capacity of metabolites (TCM) represents the overall capacity of the transport of all intermediate metabolites of one consortium. We found that the values of AMB, AFC, and TCM of the consortia positively correlated with their FRs (Figure S6), with the degradation rates increasing with increasing FR (Figure 3A). Multivariate linear regression analysis further suggested that these three parameters significantly co-determined the degradation rates of the consortia (Figure 3B; R^2^=0.24). While AMB showed a negative effect on the degradation rates (coefficient: -0.44), both AFC and TCM exhibited positive effects (coefficients: 0.31 and 0.44), which was consistent with our hypothesis. We obtained similar results in the analysis based on the growth rates of the consortia (Figure S7). Together, these findings provided indirect evidence that MCFRs possess higher functions as their configurations balanced the conflicts among the metabolic burden of strains, average functional capacity per biomass unit, as well as their capacity to transport metabolites.

**Figure 3.**
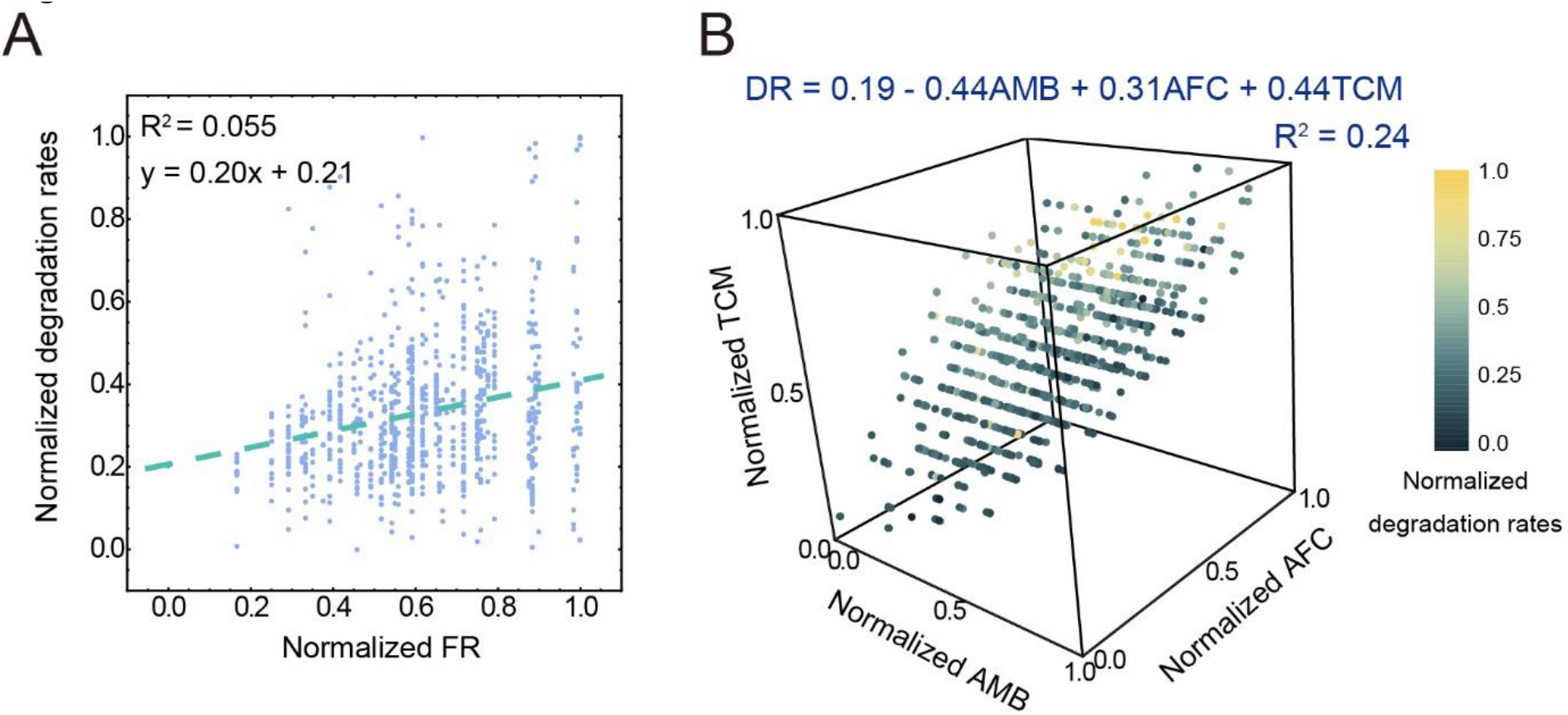
The functions of the engineered consortia can be predicted by the average metabolic burden of strains (AMB), average functional capacity (AFC), and transport capacity of metabolites (TCM). (A) The correlations between the degradation rates of the consortia and their functional redundancy level (FR). The correlation is fitted alongside linear curves (shown by the green dashed lines). (B) The summarized correlations among AMB, AFC, TCM, and degradation rates of the consortia. While the values of AMB, AFC, and TCM are shown by the three axes, the naphthalene degradation rates are shown by the color intensity. A fitted linear function given at the top of the graph indicates the fitted correlation. The definitions of these indexes are described in Supplementary Information S1.3. and Values were normalized according to the maximal and minimal values.

### Mathematical modeling extends our experimental observations and provides simple strategies to screen synthetic consortia optimal for any given pathway

To extend our observations to more pathway conditions and define an approach to screen the MCFRs possessing the best function at different pathway conditions, we next adopted a mathematical modeling strategy. We incorporated the assumptions of functional redundancy into our previously established model that characterizes a rigorous MDOL system^20^ (Methods; Supplementary Information S1.4). In line with our experimental system, we considered that a substrate can be degraded by a pathway that segregates into four steps. We assumed that the degradation was carried out cooperatively by consortia composed of four populations that could perform one, two, or three steps. We modeled the dynamics of 860 possible four-member MCFRs, one rigorous MDOL consortium (Figure S1B), and an autonomous population using a series of ordinary differential equations (Methods; Supplementary Information S1.4). The main parameters of the equations include the relative reaction rate constants of the last three steps to the first (*α_i_=a_i_*/*a_1_*, *i* = 2 – 4), the concentrations of enzymes (*e_i_*, *i* = 1 – 4), the coefficients of the metabolic burdens of producing the four enzymes (*c_i_*, *i* = 1 – 4), and the transport rates of the substrate, intermediates, and final product across the cell membrane (*γ_j_*, *j* = 1 – 5). We measured the degradation rates and growth rates to quantitatively compare the functions of the 862 systems as the set-up in our experiments (Methods; Supplementary Information S1.4).

When we used the parameter set derived from our experimental system (Table S1) for our computational simulations, we found that our model accurately predicted the degradation rates (Figure 4A) and growth rates (Figure 4B) of the MCFRs, autonomous population, and rigorous MDOL derived from our experiments. We also found that the degradation rates and growth rates of the modeled consortia increased with their levels of functional redundancy (FR; Figure S8A-B). Our multivariate linear regression analysis strongly suggested that the average metabolic burden (AMB), the average functional capacity (AFC), and the transport capacity of metabolites (TCM) co-affected the function of the consortia (Figure S8C-D). Together, these results reproduced the observations from our experiments.

**Figure 4.**
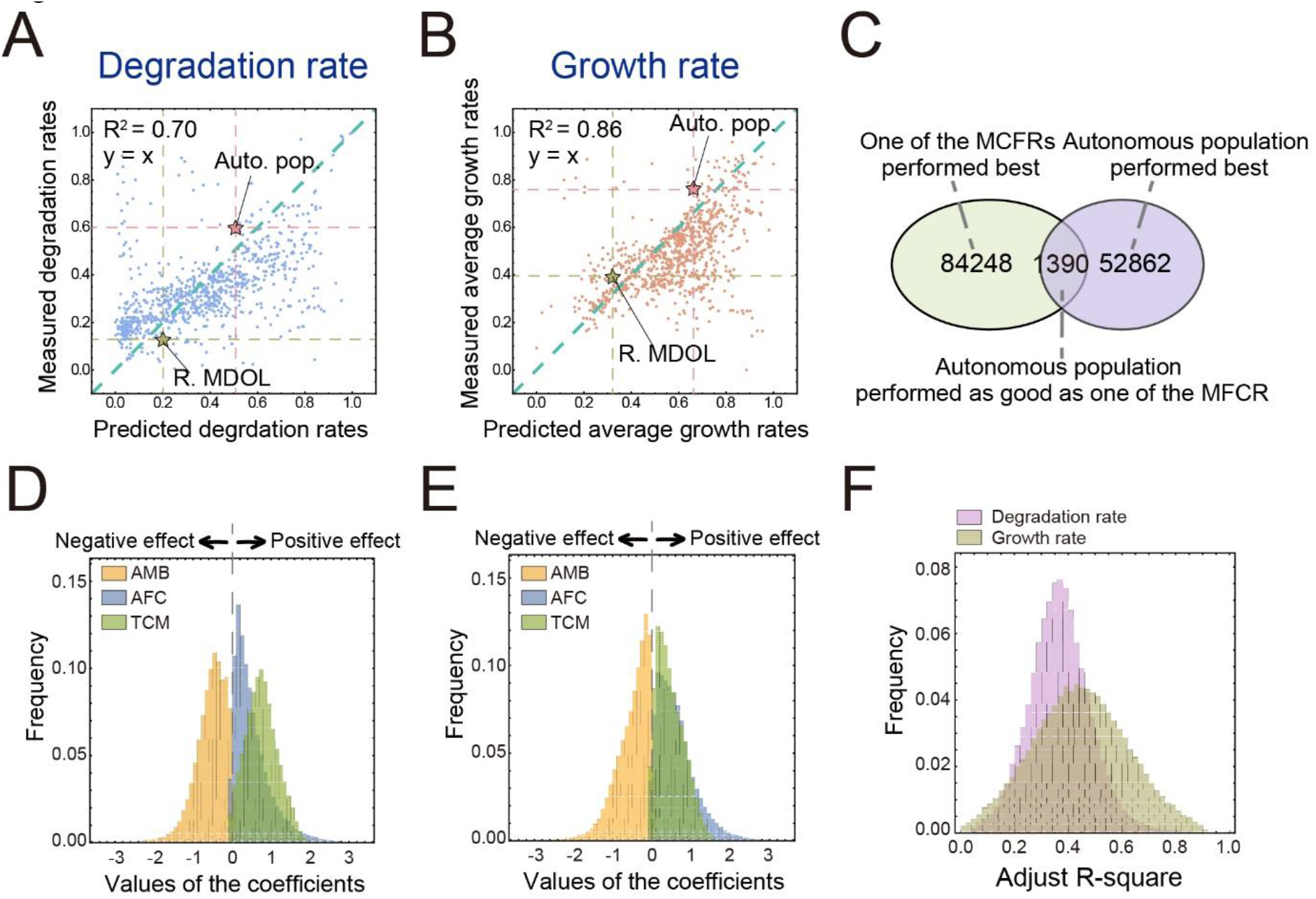
Mathematical modeling generalizes our experimental observations. (A-B) The linear correlation between the results of the mathematical simulations initialized with the parameters derived from our experimental systems and those of the wet experiments. The correlation between the simulated and experimentally measured degradation rates (A) and growth rates (B) was analyzed. The green dashed line shows the linear curve in which the predicted results are completely identical to the simulated results. The predicted and measured rates of the autonomous population (denoted by Auto. pop.; Pink) and the rigorous metabolic division of labor (R. MDOL; Yellow) are indicated by dash lines and Pentagram. (C) A Venn diagram indicates the classification of the results of the 138500 simulations into three categories. (D) distributions of correlation coefficients among the average metabolic burden of strains (AMB), the average functional capacity (AFC), the transport capacity of metabolites (TCM), and degradation rates of the consortia (E) the correlation coefficients among AMB, AFC, TCM, and the growth rates of the consortia and their FR. A coefficient value over 0 suggests the corresponding factor has a positive effect on the degradation rates or growth rates of the consortia, while a coefficient value lower than 0 suggests a negative effect. (F) the distribution of adjusted R-squared values derived from the linear regression in (D) and (E). The distributions were derived by performing fitting analysis on simulation data using 135,000 parameter sets.

#### Generalize the experimental observations in more pathway conditions

To investigate whether MCFRs are able to perform more efficiently in a variety of pathways, we conducted computational simulations using 138,500 randomly generated parameter sets (Methods; Supplementary Information S1.4.4) mimicking the deconstruction of 138,500 different metabolic pathways. We found that one of the MCFRs exhibited a higher degradation rate than the autonomous population in the simulations using 84248 (60.8 %) parameter sets (Figure 4C). When we analyzed the growth rates of the simulated microbial systems, we found that the proportion that one of the MCFRs performed better for a given parameter set was 66.9 % (Figure S9A). This result suggested that MCFRs can be more efficient systems to perform the majority of degradation pathways than the autonomous population and rigorous MDOL.

We next explored the impact of FR on the function of consortia, as well as the potential role of AMB, AFC, and TCM in determining their function under diverse pathway conditions. Our analysis showed that FR was positively correlated with the degradation rates in 95.0 % of tested conditions (Figure S10A) and with growth rates in 89.0 % of tested conditions (Figure S10B). Our data also showed that AMB affected function mostly negatively, while both AFC and TCM exhibited mostly positive effects on the function of the consortia (Figure 4D-F). This pattern was observed in 92.9 % (in terms of the degradation rate) and 90.0 % (in terms of the growth rate) of the tested conditions. Together, these results strengthened our experimental findings, suggesting that these factors co-affect the function of the consortia in a consistent manner across diverse pathway conditions.

#### Simple strategies to screen the optimal synthetic consortia for a given pathway

We next explored the strategies to select the MCFR exhibiting the highest function among all the 860 possible MCFRs under different pathway conditions. First, we analyzed the frequencies of different MCFRs that exhibited the highest degradation rates under varying parameter settings. We found that only 65 out of the 860 MCFRs achieved the highest substrate degradation rate under the condition of at least one parameter set (Figure 5A-B). We also found that the MCFR [1, 0, 1, 1], [1, 1, 0, 0], [1, 1, 0, 1] & [1, 1, 1, 0] possess the highest frequency (12.62 %) to exhibit the highest degradation rate (Figure 5A). Our analysis further indicated that the total frequency of the top 30 MCFRs that were most possible to have the highest rate reaches 95 % (Figure 5C). We observed similar trends in the analysis based on the growth rates (Figure S9B-C). These results immediately suggested a simple strategy for selecting MCFRs of the highest function: instead of measuring all the 860 consortia, the MCFR with the best function can simply be determined by measuring the functions of these top 30 MCFRs (Figure 5A: MCFRs labeled red; Figure 5B); the MCFR with the highest function among the 30 consortia should exhibit a probability of 95 % to be the consortium with the highest function of all the 860 consortia. This strategy was applicable to our experiment system, as the consortium composed of [1, 0, 1, 1] & [1, 1, 0, 0] & [1, 1, 0, 1] & [1, 1, 1, 0] was one of the 30 consortia and performed best in our experiments.

**Figure 5.**
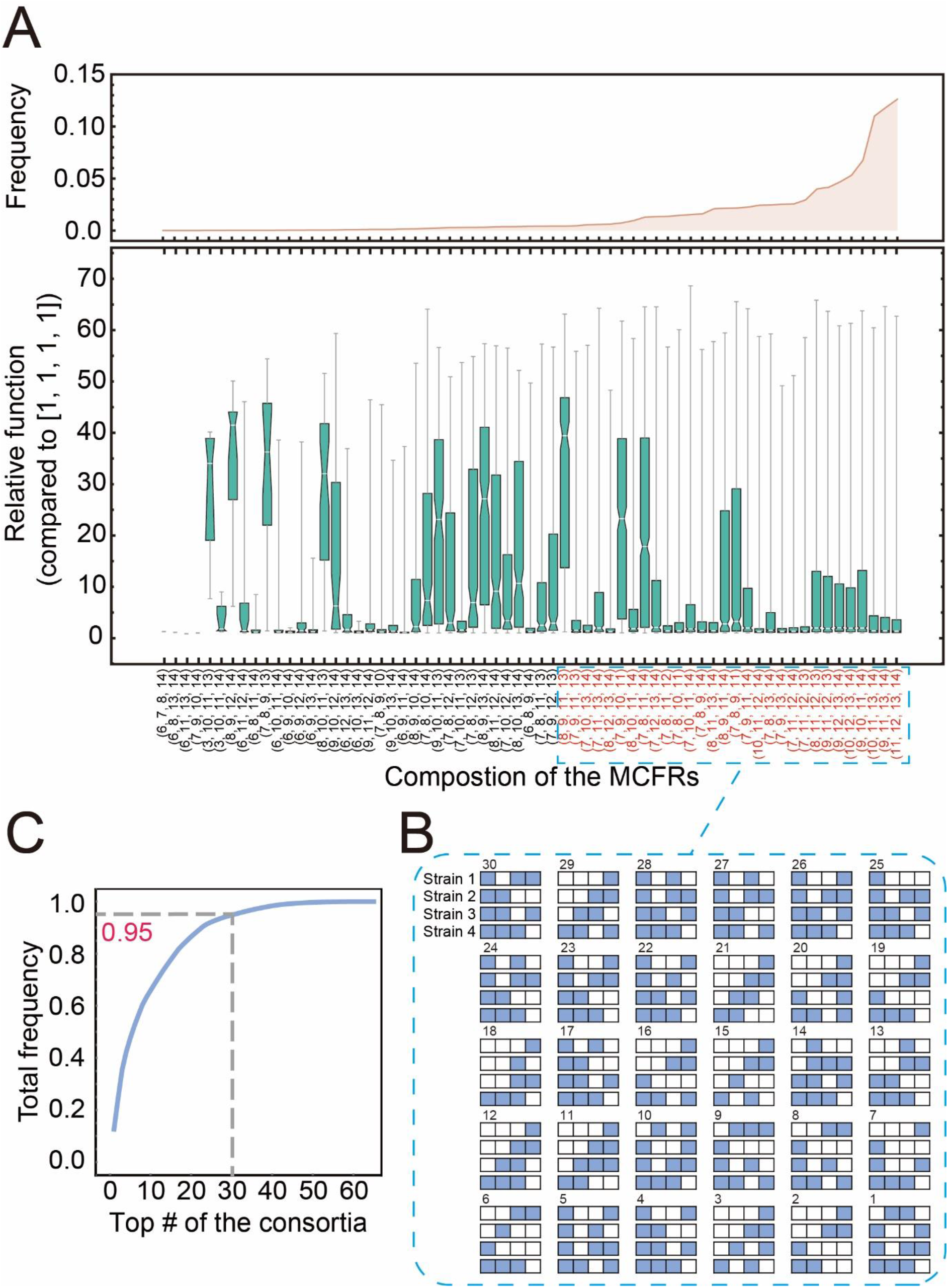
Mathematical modeling provides simple strategies to screen the optimal synthetic consortia for a given pathway. (A) Summary of the 65 consortia that possess the highest degradation rate in all simulations. The upper graph shows the frequency of each consortium possessing the highest degradation rate in the simulations. The bottom graph shows the relative degradation rates of each consortium to that of the autonomous population in all the simulations in which it possesses the highest degradation rate. The consortia are named after the four-number vectors following the same rule as in Figure 2. The names of the consortia with the 30 highest degradation rates were labeled red. (B) Diagrams of the composition of the consortia possessing the 30 highest degradation rates. The composition of each consortium is indicated by a four-row array. Each row represents the genotype of one strain involved in the consortium to perform the four-step, where the blue grid indicates that the strain possesses can perform the corresponding step while the white grid indicates it is unable to perform that step. The rank order of each consortium was denoted at the up left of each array. (C) The accumulated frequency of the consortia with the *N* highest degradation rates against the value of *N*. When the consortia with the 30 highest degradation rates are included, their total frequency equals 95 % (indicated by the grey dashed line).

Next, we performed correlation analysis to search for the most important factors that determine the function of MCFRs. As shown in Figure 6A and Figure S11A, the modeled MCFRs tended to exhibit higher function than the autonomous population when the relative reaction rates of the last three steps to the first (*α_i_=a_i_*/*a_1_*, *i* = 2 – 4), the concentrations of the last three enzymes (*e_i_*, *i* = 2 – 4), and the burdens of producing the last three enzymes (*c_i_*, *i* = 2 – 4) are low. Critically, high-function MCFRs also required a high concentration of the first enzyme (*e_1_*) and a high cost of producing the first enzyme (*c_1_*). Next, we explored how these parameters determine which MCFR possesses the best function. Our Point Biserial Correlation analysis indicated that seven parameters, namely *α_i_* (*i* = 2 – 4) and *γ_j_* (*j* = 2 – 5), were the key parameters that determine the MCFR possessing the best function (Figure 6B; Figure S11B; Figure S12). For example, if the values of *α_2_* and *γ_2_* for a given scenario were low (i.e., *α_2_* falls into the smaller 43.75% of our tested range and *γ_2_* falls into the smaller 18.75 % of its range; Figure S12A), and the values of *α_3_*, *α_4_*, and *γ_3_* were high (i.e., *α_3_* and *α_4_* fall into the larger 50% of our tested range and *γ _2_* falls into the larger 68.75 % of its range; Figure S12A), the MCFR composed of [1, 0, 1, 1] & [1, 1, 0, 0] & [1, 1, 0, 1] & [1, 1, 1, 0] exhibited the best function. Together, these results suggest that it is feasible to select the MCFR exhibiting the highest function on the basis of the measured values of the seven key parameters.

**Figure 6.**
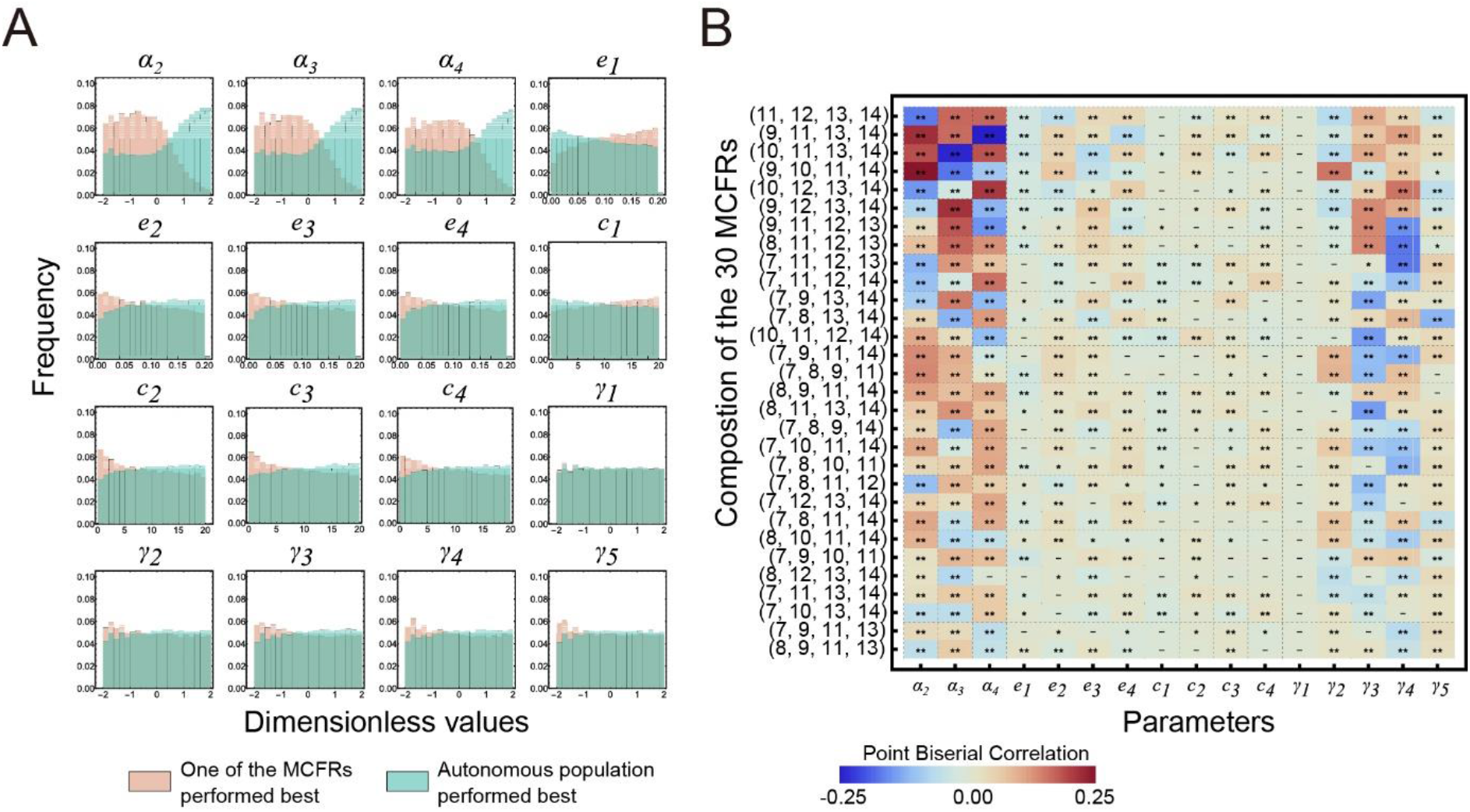
Key parameters that determine the function of consortia performing metabolic division of labor and possessing functional redundancy (MCFRs). (A) Distributions of two groups of parameter values lead to two different types of simulation results. The light red histograms represent values of the 84248 sets resulting in the simulations in which one of the MCFRs possesses the highest degradation rate. The green histograms represent values of the 52862 sets resulting in the simulations where the autonomous population performs better than all the consortia. (B) Point Biserial Correlation suggests how the values of different parameters determine whether an MCFR becomes the consortium with the highest degradation rate. We set a value of 1 if an MCFR possesses the highest degradation rate in a simulation initialized with a parameter set and a value of 0 if it does not perform best. As a result, a binary variable is obtained. Then Point Biserial Correlations between the values of each parameter and this binary variable are performed. The values of the derived coefficients are shown by the color intensity. Here, the results of the 30 consortia with the highest frequencies to possess the highest degradation rates were shown. The names of these consortia are presented following the same rule as in Figure 2. The markers involved in each grid are derived from Mann-Whitney Tests between the set of the parameters that leads to the corresponding MCFR possessing the highest degradation rate and that leads to MCFR does not perform best. “**”: *p* < 0.0001; “*”: *p* < 0.01; “-”: *p* > 0.01.

### Species exhibiting metabolic versatility while being non-autonomous are prevalent in natural communities

Based on these findings, we hypothesized that many microorganisms evolve genotypes that can perform one or multiple steps, but not all steps, of a particular metabolic pathway. These microorganisms may form MCFRs exhibiting different configurations, which potentially improve pathway efficiency. We tested this hypothesis in the pathways for the degradation of eight typical hydrocarbons, including three aliphatic (short-chain and long-chain n-alkanes, as well as cycloalkane) and five aromatic hydrocarbons (toluene, phenol, xylene, benzene, biphenyl, and naphthalene). We divided these pathways into several steps according to the following two conditions: (1) whether the selected intermediates are chemically stable and (2) whether the intermediates can be transported across the cell membrane so that be exchanged among different populations (Figure S13). To identify the distribution of the genes encoding the enzymes responsible for every metabolic step in microbial genomes, we searched for the hydrocarbon degradation genes in a database recently built from 24,692 publicly available archaeal (*n* = 1246) and bacterial (*n* = 23,446) genomes^21^. Based on these analyses, we classified these microorganisms into different genotypes that can perform one or multiple metabolic steps, which are conceptualized by bit strings containing “0”, “1”, “A” and “B” (Figure 7; Methods). The definition of ‘0’ and ‘1’ are identical throughout this report. For a metabolic step possessing multiple shunt reactions, we used “A” or “B” to denote that one genotype harbors the genes to finish this step via different shunt reactions. For example, genotype [A, 0, 0] in toluene degradation indicates that the genotype can only perform the first step of toluene degradation via the shunt reaction “A” (toluene-2,3-deoxygenation) but cannot perform the remaining two steps (Figure 6E; Figure S13E). These definitions allowed us to analyze the frequency of different genotypes contributing to each pathway.

**Figure 7.**
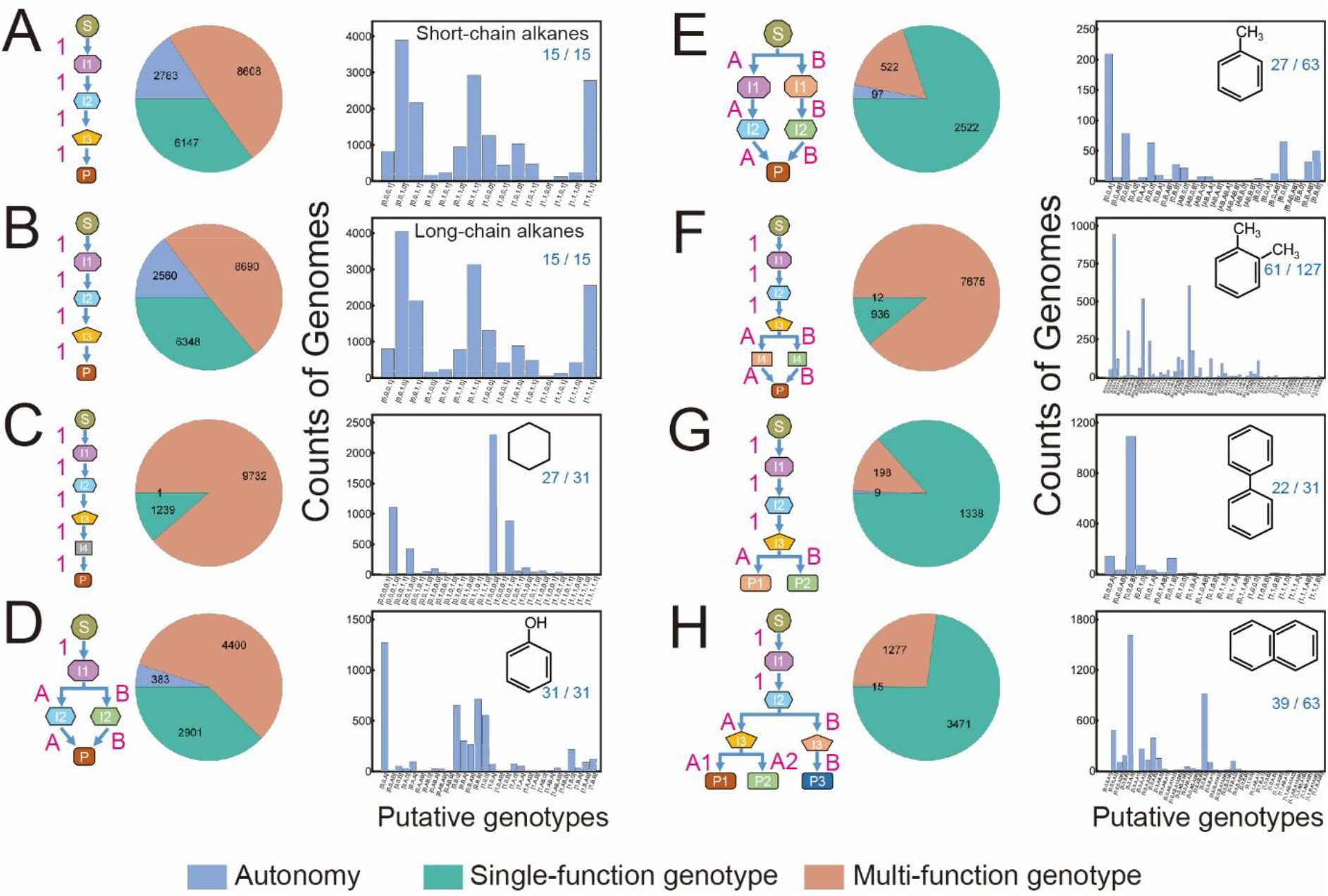
Distributions of hydrocarbon-degrading genes in microbial genomes. Genes involved in the degradation pathways of eight hydrocarbons were analyzed, including short-chain n-alkanes (A), long-chain n-alkanes (B), cycloalkane (C), toluene (D), phenol (E), xylene (F), benzene (G), biphenyl (H), and naphthalene (I). In each graph, the left diagram shows how the degradation pathway was deconstructed according to (1) whether the selected intermediates are chemically stable and (2) whether the intermediates can be transported across the cell membrane so that be exchanged among different populations. “S” indicates the initial substrate; “I” indicates different intermediates; “P” indicates the final product. Details of these pathways are shown in Figure S12. The middle pie chart shows the frequencies of the putative autonomous genotype that can autonomously perform the pathway (the blue sector), the putative genotypes that can only perform one metabolic step (the green sector), and the putative genotypes that can perform multiple but not all steps (the light red sector). The right bar chart shows the frequencies of all the putative genotypes. These genotypes were conceptualized by bit strings containing “0”, “1”, “A” and “B”. For a metabolic step that only has one known reaction, “1” denotes that the genotype can perform this reaction while “0” denotes that it is unable to perform it. For a metabolic step that possesses multiple shunt reactions, “A” or “B” denotes that the genotype has the genes to execute this step via “A” or “B” shunt reactions; “0” denotes that the genotype is unable to perform this step regardless which shunt reaction.

We found that only a small percentage of microbes (0.009 % ∼ 15.9 %) is present as an autonomous population that contains all enzymes required to completely degrade one hydrocarbon compound, especially in the pathways mediating the degradation of aromatic hydrocarbons (0.009 % ∼ 5.0 %; Figure 6). In contrast, most microbes lack a complete set of enzymes. Instead, many microbes contain a subset for hydrocarbon degradation, an observation that is in agreement with findings in several previous studies^21-24^, suggesting that MDOL is prevalent in hydrocarbon degradation. When we assessed three degrading pathways of aromatic hydrocarbons, we found that most microbes only perform one single metabolic step (toluene: 80.3 %, Figure 7E; biphenyl: 86.6 %, Figure 7G; naphthalene: 72.9 %, Figure 7H). In the other five degrading pathways, we found that genotypes with the ability to perform multiple steps of the pathways were present at a higher abundance (short-chain n-alkanes:

49.1 %, Figure 7A; Long-chain n-alkanes: 49.4 %, Figure 7B; cycloalkane: 88.7 %, Figure 7C; phenol: 57.3 %, Figure 7D; xylene: 89.0 %, Figure 7F). This result supports our hypothesis that many microorganisms evolve to be non-autonomous genotypes but maintain abilities to perform one or multiple steps of metabolic pathways. These microorganisms presumably form MCFRs for the highly efficient degradation of hydrocarbons, a process that has been reported for several natural microbial communities^22-24^.

## Discussion

Here, we demonstrated that introducing functional redundancy to a consortium performing metabolic division of labor (MDOL) generates a novel consortium possessing a higher function. Our mathematical modeling provides general strategies to identify the MDOL consortium with functional redundancy (MCFR) that exhibit the best function among a series of possible MCFRs.

Synthetic ecology represents a new frontier for synthetic biology. Through the design of synthetic consortia that are composed of multiple interacting microbes, it can address a number of issues that cannot be addressed in single microbes^25-28^. MDOL represents a commonly-used strategy for engineering synthetic consortia^12,13,29,30^ that are applied for the degradation of environmental pollutants^20,31-33^, the production of valuable products, such as biopolymers^34,35^ and pharmaceuticals^36,37^, as well as for the generation of biofuels^38-41^. However, most MDOL consortia are designed as rigorous MDOL systems, which distribute different metabolic tasks among different specialists, each of which only performs one specific task. A small number of recent studies attempted the engineering of MDOL consortia by allowing one strain to perform multiple metabolic tasks, thus increasing functional redundancy at the community level. For example, one recent study^42^ on the degradation of atrazine assembled four bacterial strains into a consortium, in which three strains redundantly performed the upper pathway (degrading atrazine into cyanuric acid), with two other strains performing the lower pathway (further degrading cyanuric acid). The consortium achieved high overall activity of atrazine mineralization. In another study that aimed at lignocellulose biotransformation, the community was composed of three strains, in which one strain produced the enzyme for cellulolytic activity, and the other two strains redundantly provide the enzymes for ligninolytic activity^43^. Our work here offers a systemic assessment of the functions of different MCFRs, indicating that introducing functional redundancy into MDOL consortia is highly probable to achieve a higher community function. Our mathematical modeling presented here further suggests that this strategy is feasible to increase the ability of synthetic consortia to perform the majority of degrading pathways. Our model also offers simple strategies for selecting the MCFR exhibiting the best function. We anticipate that our findings will provide ample guidance for further studies investigating pathway optimization in synthetic consortia.

The functional redundancy within a community considerably affects its ecological properties^18,44-46^. Previous theories mainly focused on the relationship between redundancy and stability^44,45,47,48^, while few studies focused on the relationship between functional redundancy and ecosystem functioning. While one study showed that functioning is consistently increased with redundancy^49^, another study showed that functioning remains independent of redundancy levels^50^. Our data suggest a positive redundancy-functioning relationship, in agreement with the former opinion. Importantly, our analysis offers a more mechanistic understanding of this relationship and shows that increasing redundancy has both positive and negative effects on ecosystem functioning. While increasing redundancy enhances the functional capacity and the transport of metabolites that benefit community functioning, it also imposes a high metabolic burden that harms the proper community function. Therefore, the “optimal” state of functional redundancy that enables the highest ecosystem functioning emerges after the trade-off of these detail factors. This knowledge may help to artificially control and regulate the composition of microbial communities toward optimal performance.

Our findings also offer an explanation for why natural communities tend to exhibit a prevalence of functional redundancy. The classical *Competition Exclusion Theory* asserts that two species occupying identical niches cannot coexist in a well-mixed system^51^. This prediction is opposed to the idea of the prevalence of high functional redundancy concerning metabolic pathways in natural communities^18,52^ if functional redundancy is defined as the coexistence of organisms that share the exact same set of functions (termed “strict redundancy” in the reference^52^).

Several recent studies proposed that functional redundancy may be promoted by differentiation together with other niche axes^18,52^. In such a scenario, organisms only share a subset of specific functions but differ in other functions or other ecological requirements (termed “partial redundancy” in the reference^52^). The design of the consortia used in our study is based on the scenario of “partial redundancy”. For example, strains [0, 0, 1, 1] and [0, 1, 0, 1] share the function of performing the last step but differ in the function of performing the second and third steps. Our analysis of the community structures supports the idea that ‘partial redundancy’ can be stably maintained in communities (Figure S14). Our genomic analysis suggests that such a form of “partial redundancy” is prevalent in the community performing hydrocarbon degradation. Previous metagenomic studies also observed similar phenomena in the degradation of other organic compounds^42^, as well as other metabolic activities, such as sulfur oxidation, denitrification, and sulfate reduction^53^. As an optimal level of redundancy can enhance the function of a community, it can be assumed that those communities which maintain an adequate level of functional redundancy are likely to be favored selectively in nature, especially when environmental pressures act on communities with higher function (for instance, a community-consuming substrate and accumulating biomass at a faster rate). To further test this hypothesis, the large-scale metagenomic analysis should be used for analyzing the distributions of functional genes of different microorganisms within the same habitat.

Our results also provide novel insights into how MDOL communities assemble. Our genetic analysis shows that the genotypes that only performed the downstream pathway (the last two steps) are characterized by a higher abundance than the genotypes that performed one or two steps of the upstream pathway (Figure S14). This result suggests that microbes possessing the downstream pathways may be selectively favored in the presence of MDOL in natural communities. We also obtained similar results in our genomic investigation. For instance, those single-function microbes usually possess the exclusive ability to perform the last step of the pathway (Figure 7E). Specifically, most microbes only maintain genes encoding catechol degradation, the shared last step for aromatic hydrocarbon degradation (Figure 7; Figure S13). These observations agree with the findings in our recent studies^20,54^, which show that the strain performing the last step has preferential access to degradation products. Since these products are the carbon sources that support strain growth, the strains performing the downstream pathways may gain more benefits than others and thus will be selectively favored. In agreement with our findings in our previous report, this study again indicates that benefits unevenly allocated between different members critically affect the assembly of a community, and ultimately interfere with its proper performance.

While our study provides valuable insights into the assembly and function of MDOL communities, it is important to note one limitation. The rules and strategies developed in the study are primarily based on scenarios of the division of labor on the degradation pathways of organic compounds, where substrate degradation provides available carbon sources to support the growth of different populations. As a result, our findings may not be applicable to the engineering of specific consortia established for other purposes. For instance, in synthetic consortia designed to synthesize targeted products, populations are typically supplied with sufficient carbon sources to support their growth^13,36,37^, which requires different assumptions to link the division of labor in the pathways and the growth of the populations. A more comprehensive understanding of these factors can aid in the development of more effective and efficient microbial systems.

In summary, our study demonstrates that functional redundancy is a nonnegligible factor determining the function of a microbial system. We propose a general strategy guiding to control of this factor and thus provide novel insight into the *de novo* designing and engineering of high-performance microbial systems. Our results also raise a novel explanation for how functional redundancy is maintained and play a role in natural communities, increasing our understanding of how natural communities assemble and evolve in their perpetual struggle for survival within their ecological niche.

## Methods

### Construction of the strains involved in the synthetic microbial consortia

The strains and plasmids used in this study are summarized in Table S3. All strains used in our synthetic consortia were engineered from a naphthalene-degrading bacterial strain *P. stutzeri* AN10^55,56^. The engineering workflow is summarized in Figure S2. Briefly, an autonomous strain was engineered by removing the *nahW* gene (encoding a salicylate hydroxylase) and the *catA* gene (encoding a catechol 1, 2-dioxygenase) of *P. stutzeri* AN10. In addition, the original promoters of the two naphthalene degradation operons (induced by one of the intermediates, salicylate^57,58^) with an IPTG-induced promoter, P ^59,60^. The derived autonomous strain, namely [1, 1, 1, 1], degrade naphthalene autonomously, and its genes encoding the enzymes for naphthalene degradation are all located in the two engineered operons. The naphthalene degradation pathway in [1, 1, 1, 1] was further partitioned into four metabolic steps. The four key enzymes responsible for these four steps were encoded by *nahA* gene (encodes a naphthalene dioxygenase), *nahC* gene (encodes a 1, 2-dihydroxynaphthalene dioxygenase), *nahG* gene (encodes a salicylate hydroxylase), and *nahH* gene (encodes a catechol 2, 3-dioxygenase). To construct the mutants that can only perform one or a subset of metabolic steps in naphthalene degradation, the four key genes were knocked out one by one following the standard workflow shown in Figure S2. As a result, forty mutants were obtained. These strains were named using bit-strings, in which ‘1’ denotes that the related key gene is retained in the strain, thus it can carry out the corresponding step. In contrast, ‘0’ denotes that the related key gene is defective, thus it is not capable of carrying out the corresponding step. All these genetic manipulations were implemented by allele exchange using the suicide plasmid pK18mobsacB^61,62^. The constructed strains were validated by PCR and DNA sequencing. In addition, enzymic activity assays and monoculture experiments were performed to verify the phenotypes of different strains, following the methods reported before^63-65^. More details of strain construction and verification are provided in Supplementary Information S1.1.

### Construction and culturing of the synthetic microbial consortia

In total, 1456 synthetic microbial consortia were constructed by grouping two, three, or four engineered strains. The strains were cultured in 25-mL flasks containing 5 mL of new fresh medium at 30°C by shaking at 220 rpm. To prepare the inoculum, *P. stutzeri* strains were first grown at RB liquid medium (Yeast extract 10 g/L, beef extract 6 g/L, peptone 10 g/L, ammonium sulfate 5 g/L). The cells were then washed twice with the minimum medium^66^ to make an inoculum. For co-culture experiments, the inocula of the two, three, or four strains involved in a synthetic consortium were concentrated to an Optical Density (OD, measured at 600 nm) of 5.0, and mixed at equal initial abundance, or a pre-designed initial ratio. The cultures were then inoculated to a 25-mL flask containing 5 mL new fresh minimum medium (starting OD: 0.05) supplemented with naphthalene powder (1% w/v) as the sole carbon source and 1 mM IPTG (to induce the expression of naphthalene degradation genes). Six replicates were performed for each consortium, three of which were used for growth measurements. The other three replicates were used for the measurements of naphthalene degradation rates. To quantify the growth rates of each consortium, 100 μL of culture liquid was regularly sampled from the medium during a culture period of 144 hours for OD measurements. The derived growth curves were fitted to Logistic Function^67^ to calculate the growth rates.

### Measurements of naphthalene degradation rates

The residual naphthalene after culture was measured by Gas Chromatography-Mass Spectrometer (GC-MS) and then used for the quantification of the naphthalene degradation rate of the synthetic consortia. Briefly, 5 mL of the bacterial culture was mixed with 1.5 mL of *n*-hexane after 96-h. The mixture was repeatedly pipetted until all the naphthalene solid particles were completely dissolved in *n*-hexane. This mixture was then centrifuged at 10,000 g for 10 min to allow clear stratification of the organic phase (*n*-hexane) and aqueous phase. The organic phase was then collected and filtered through a 0.22 μm filter into a brown chromatography bottle. The naphthalene concentration of the sample was measured using Agilent 7890A gas chromatography paired with the Agilent 5975C mass spectrometer. The inlet temperature was set to 295℃. Helium was used as the carrier gas with a flow rate of 1 μL/min. The injection volume was 1.0 μL. The temperature program was 50℃ (1 min isothermal), 50 to 250℃ (20℃ / min), and 250℃ (3 min isothermal). For the mass spectrometry, the ion source temperature, ionization energy, interface temperature, and quadrupole temperature are set to 230°C, 70 eV, 280°C, and 150°C respectively. For determining the concentration of naphthalene, six internal standards were employed (naphthalene at 0.1 g/L, 0.5 g/L, 3 g/L, 10 g/L, 30 g/L, and 60 g/L in hexane). The obtained GC-MS data were uploaded to XCMS (https://xcmsonline.scripps.edu/), an open-source online analysis platform, to quantify the peak area of naphthalene. To obtain the residual amount of naphthalene in each sample, the data were compared to the standard curve plotted using the internal standards. Finally, the naphthalene degradation rate (NDR) is calculated as follows:

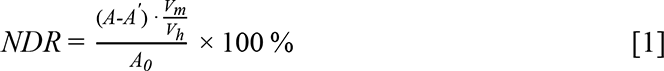

Here, *A* is the residual amount of naphthalene in the sample; *A^’^* is the residual amount of naphthalene in a control sample without the inoculation of bacteria; *A_0_* is the initial amount of naphthalene; *V_m_*is the volume of n-hexane for naphthalene extraction; *V_h_* is the volume of the culture medium.

### Quantifying the relative frequencies of different strains in our synthetic consortia

To qualify the relative frequencies of different strains in our consortia, we knocked in specific barcodes to the chromosomes of each strain. Then the relative frequencies of different strains are determined by high-throughput amplicon sequencing targeted on those barcode regions. The detailed protocols and the defense of the methodology were described in Supplementary Information S1.2.

### Formulation of the Mathematical model and the simulation protocols

The mathematical model was modified from the model of our previous studies that characterize the scenario of the rigorous metabolic division of labor (MDOL)^20^. The detailed derivations of the model are described in Supplementary Information S1.4. The definitions and dimensionless methods of all the variables and parameters are listed in Table S1-S2. In this model, the dynamics of four-member MDOL consortia possessing functional redundancy (MCFRs) were characterized. Identical to our experimental system (Figure 1 and Figure S1), a degradation pathway was divided into four steps. One initial substrate (S), three intermediates (I1, I2, I3), and an end product (P) were included. One modeled consortium was composed of four strains, each of which performed one, two, or three metabolic step(s) of the pathway. The genotypes of the strains were conceptualized by bit strings containing “0” and “1” (*ɛ_k_*). Ordinary differential equations (ODEs) were used to formulate the dynamics of intracellular and extracellular intermediates and end products, as well as the growth of all the strains involved in the community. In all cases, the models were built on a well-mixed system. For simplicity, the model was built based on seven simple assumptions identical to our previous study^20^, namely transport via passive diffusion, intracellular metabolic reactions, negligible abiotic degradation of I and P, excess of initial substrate, as well as low levels of intracellular accumulation of I and P; importantly, P was assumed to be the exclusive as well as a limited resource for the growth. Here, the dimensionless forms of the models are presented:

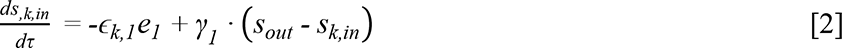

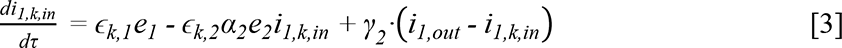

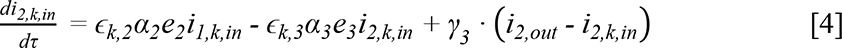

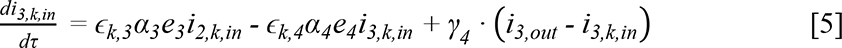

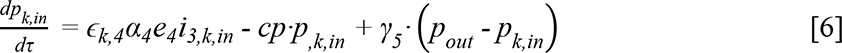

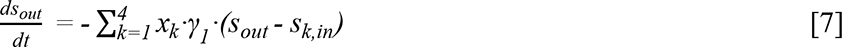

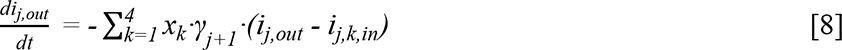

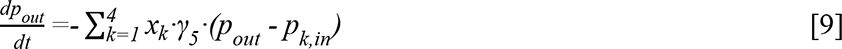

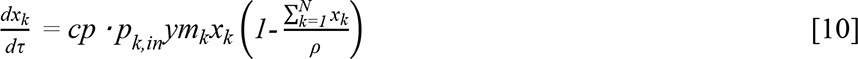

In the model, *j* = 1 ∼ 3 (i.e., three intermediates) and *k* = 1 ∼ 4 (i.e., four genotypes involved in the consortium). *s_k,in_* represents the intracellular S concentration of the *k*th strain; *s_out_* represents the extracellular concentration of S; *i_j,k,in_*represents the intracellular concentration of the *j*th intermediate of the *k*th strain; *i_j,out_*represents the extracellular concentration of the *j*th intermediate; *p_k,in_* represents the intracellular P concentration of the *k*th strain; *p_out_* represents the extracellular concentration of P; *ɛ_k_* represents a vector that characterizes the genotype of the *k*th strain; *x_k_* is the biomass of the *k*th strain; *a_k_* is the reaction rates of the *k*th reactions; *γ_j_* (*j* = 1 ∼ 5) are the diffusion rates of S, I1, I2, I3, and P across the cell membrane; *cp* are the consumption rate of P of the strains; *ρ* is the carrying capacity of the whole communities; *y_k_* is the yield coefficient for biomass production of the *k*th strain; *m_k_*is the metabolic burden of the *k*th strain.

Details of the simulation and analysis protocols of our model and the downstream analyses are described in Supporting Information: S1.4. Briefly, numerical simulations of the model were performed using the NDsolve function of Wolfram Mathematica (version 12.0). The numerical solutions of all variables, including the dynamics of mass (S, I, P) concentration and biomass, were recorded for further analyses. To mathematically predict the experimental results, parameter values matching our experimental system (Table S1) were applied to the model and 862 systems (meaning different combinations of *ɛ_k_*) were simulated. To generalize the experimental findings in more pathway conditions, 138500 parameter sets were generated by randomly picking the values of the 16 main parameters from the given ranges obtained from literature search or experimental measurements (Table S1). All the simulations, as well as the downstream analysis, were performed using custom Mathematica scripts. The source codes used for the model analysis are publicly available (https://github.com/WMXgg/MDOLcode/tree/master/MDOL-redun).

### Analysis of the distributions of hydrocarbon-degrading genes in microbial genomes

The analysis was performed based on a database compiled recently^21^, which used Annotree to annotate the genes encoding the aerobic degradation pathways of aliphatic (short-chain and long-chain n-alkanes, as well as cycloalkane) and aromatic (toluene, phenol, xylene, benzene, biphenyl, and naphthalene) hydrocarbons in 24,692 genomes from 123 bacterial and 14 archaeal phyla. The database was further analyzed in this work. First, eight degradation pathways were divided into three to six steps according to two conditions: (1) whether the selected intermediates are chemically stable and (2) whether the intermediates can be transported across the cell membrane so that be exchanged among different populations (Figure S12). Next, the genes responsible for each step were searched in the database to obtain their distributions in all the genomes. Accordingly, the genotypes of all these microorganisms were determined, representing their potential for how many metabolic steps they can perform for these degradation pathways. These genotypes were conceptualized by bit strings containing “0”, “1”, “A” and “B”. For a metabolic step that only has one known reaction, “1” denotes that the genotype can perform this reaction while “0” denotes that it is unable to perform it. For a metabolic step that possesses multiple shunt reactions, “A” or “B” denotes that the genotype has the genes to execute this step via “A” or “B” shunt reactions; “0” denotes that the genotype is unable to perform this step via any shunt reactions. The distribution of the different genotypes in all eight pathways is summarized and visualized in Figure 6. All analyses were performed using custom-tailored Mathematica scripts. The source codes used for the model analysis are publicly available (https://github.com/WMXgg/MDOLcode/tree/master/MDOL-redun).

### Quantification and statistical analysis

The LinearModelFit function in *Wolfram Mathematica* (version 12.0) was used for the linear fit of the simulation or experimental data, while the NonlinearModelFit function was used for the non-linear fit, both with default settings. The values of adjusted R-squared can be found in all related figures. Student’s T-test, Point Biserial Correlations, and Mann-Whitney Test were calculated using the custom Mathematica scripts. All stats methods are also briefly described in the relevant figure legends.

### Replication and randomization

Replicate experiments have been performed for all key data obtained in this study. Biological or technical replicate samples were randomized where appropriate. The numbers of replicates are listed in the related figure legends.

## Acknowledgments

We wish to thank Dr. Min Lin (Chinese Academy of Agricultural Sciences, Beijing, P.R. China) for providing plasmid pK18mobsacB and pRK2013, used for genetic engineering in this work; Dr. T. Juelich (UCAS, Beijing) for linguistic assistance during the preparation of this manuscript. We thank the members of the Wu lab at Peking University and Microbial Systems Ecology group at ETH Zürich for the critical discussions of this work. Finally, the author, Miaoxiao Wang, would like to acknowledge his wife, Yaxi Li, for her unwavering support and endless inspiration that has been instrumental in the success of this research. Her intelligence and charm have been a constant source of motivation and encouragement throughout this project. This work was supported by National Key R&D Program of China (2018YFA0902100 and 2018YFA0902103), National Natural Science Foundation of China (91951204, 31761133006, 31770120, and 31770118), and Sino Swiss Science and Technology Cooperation (SSSTC) Program (IZLCZ0_206044).

## Author Contributions

MW and YN were involved in the conceptualization of the study. MW designed the experiments. MW, XC, YF, XZ, and TH performed the experiments. MW constructed the ODE models and performed mathematical simulations. MW analyzed the data and wrote the original draft. YN and XLW edited the manuscript. YN, MW, and XLW raised the funding for the project.

