## Supplementary Text, Tables and Figures for "The trade-off between individual metabolic specialization and versatility determines the metabolic efficiency of microbial communities"

**Supplementary Information**

**Introducing functional redundancy as a powerful strategy for pathway optimization in synthetic consortia performing metabolic division of labor**

Miaoxiao Wang^1, 2, 3^, Xiaoli Chen^1, 4^, Yuan Fang^6^, Xin Zheng^6^, Ting Huang^6^, Yong Nie^1#^ and Xiao-Lei Wu^1, 4, 5#^

^1^ College of Engineering, Peking University, Beijing 100871, China

^2^ Department of Environmental Systems Science, ETH Zürich, Zürich, Switzerland

^3^ Department of Environmental Microbiology, Eawag, Dübendorf, Switzerland

^4^ Institute of Ocean Research, Peking University, Beijing 100871, China

^5^ Institute of Ecology, Peking University, Beijing 100871, China

^6^ School of Resource and Environmental Engineering, Hefei University of Technology, Hefei 230000

^#^Corresponding author: Research Scientist, College of Engineering, Peking University.

^#^Corresponding author: Professor, College of Engineering, Peking University.

[*S1.1.1 Construction of the strain [1, 1, 1, 1] that degrades naphthalene autonomously*	3](#_Toc126780990)

### S1 Supplementary Methods

#### **S1.1** **Construction of the strains involved in the synthetic microbial consortia**

##### *S1.1.1 Construction of the strain [1, 1, 1, 1] that degrades naphthalene autonomously*

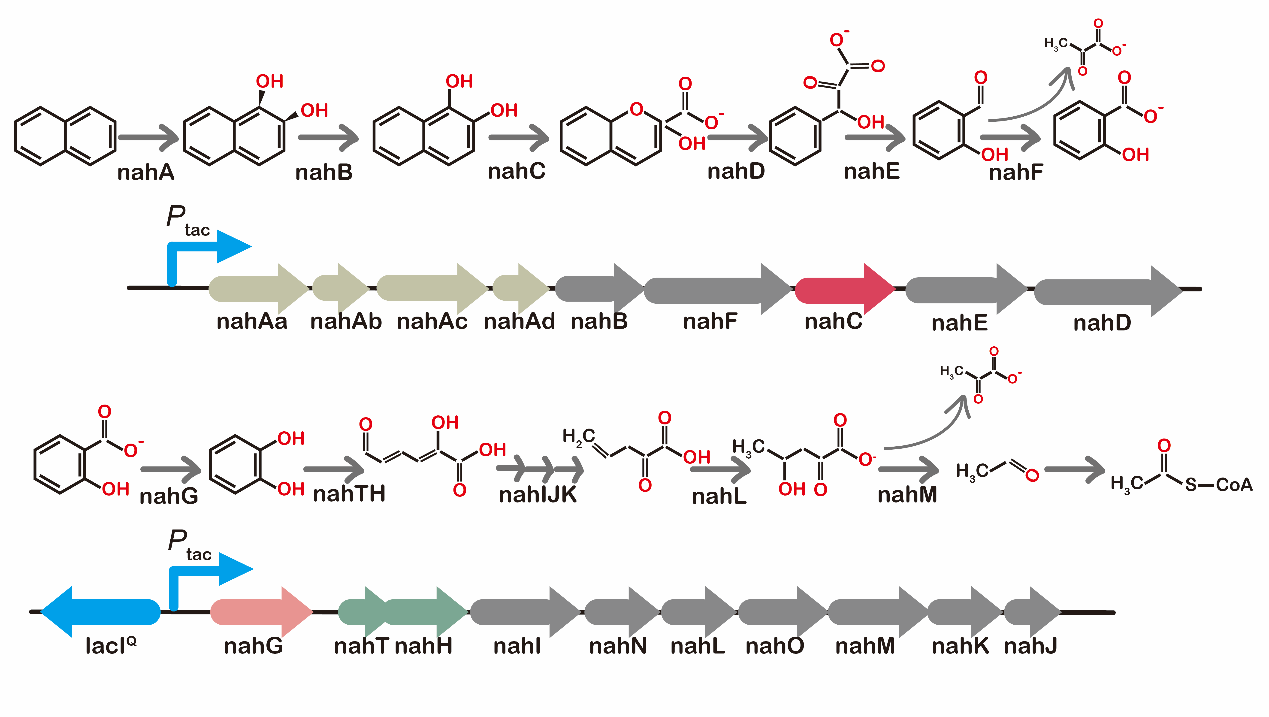

**Figure S1.1** The upstream and downstream gene clusters of naphthalene degradation in *P. stutzeri* AN1111 and their corresponding degradation pathways.

The strains used for experimental verification were all engineered from a *Pseudomonas stutzeri* strain AN10^1-3^, which possesses a great ability to degrade naphthalene autonomously through a long pathway. Our preliminary design is to knock out its genes encoding the enzymes responsible for each metabolic step, aiming to construct the mutants that are only capable of performing a single metabolic step, and then co-culture these strains to execute MDOL. In strain AN10, the genes encoding the enzymes for naphthalene degradation were reported to be mainly located in two operons^2,3^. However, when we searched its genome, we found another two groups of genetic elements that may also be included in the naphthalene degradation pathway, that is, a *nahW* gene encoding a salicylate hydroxylase^2,4,5^ and a Cat gene cluster encoding enzymes for the degradation of catechol (one of the intermediates in naphthalene degradation). Therefore, we first knocked out the *nahW* gene and the *catA* gene encoding a catechol 1,2-dioxygenase^6^. Then we replaced the original promoters of the two operons (induced by one of the intermediates, salicylate^2,3^) with the IPTG-induced promoter, *P_tac_*^7,8^. The derived strain can degrade naphthalene autonomously, and its genes encoding the enzymes for naphthalene degradation are all located in the two engineered operons (Figure S1.1). Therefore, its ability for naphthalene degradation can be quantitatively controlled by modulating the addition of IPTG (Figure S1.2). We named this controllable ‘super strain’ *P. stutzeri* AN1111 (denoted as [1, 1, 1, 1] in the main Text), denoting that it can execute all the metabolic steps of naphthalene degradation.

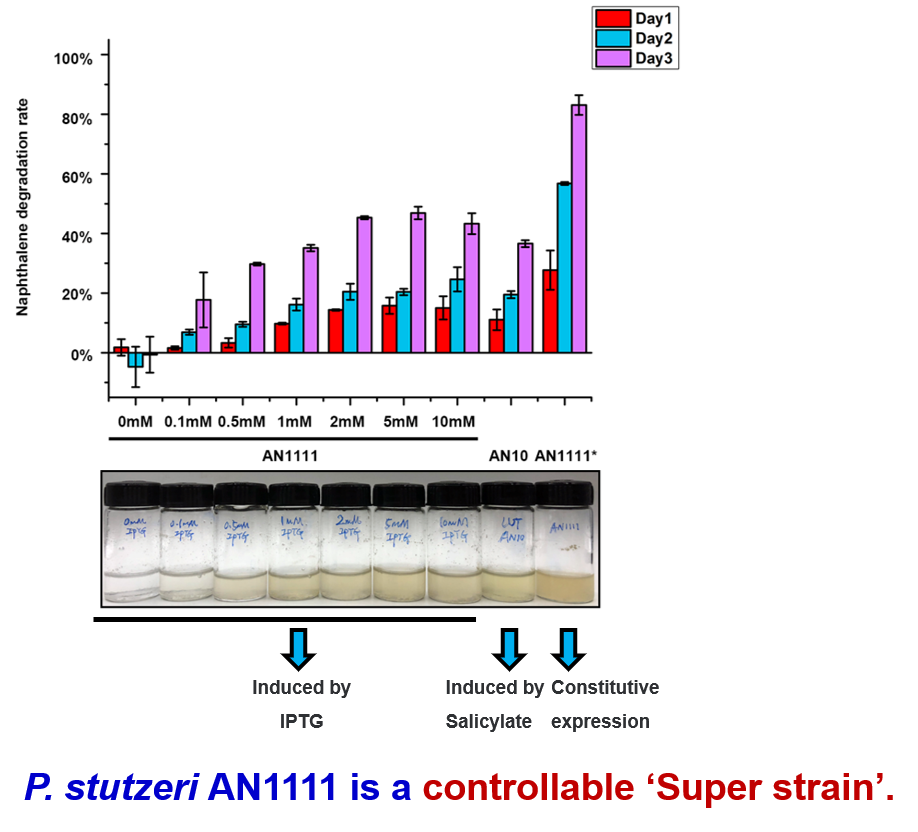

**Figure S1.2** Measurement of the naphthalene degradation rate and growth of the ‘super strain’ *P. stutzeri* AN1111 under different IPTG concentrations. Note that *P. stutzeri* AN10 is the wild-type strain, and *P. stutzeri* AN1111^*^ is a derived strain that lacks the *lacI^Q^* gene, thus the expression of naphthalene degradation genes in this strain is constitutive.

##### *S1.1.2 Construction of the mutants that are only able to perform a subset of metabolic steps of naphthalene degradation*

The naphthalene degradation pathway in *P. stutzeri* AN1111 was further divided into four steps, which were associated with three intermediates, 1, 2- hydroxynaphthalene, salicylate, and catechol (Figure S2). The four key enzymes responsible for the four metabolic steps were encoded by *nahA* gene (encodes a naphthalene dioxygenase), *nahC* gene (encodes a 1, 2-dihydroxynaphthalene dioxygenase), *nahG* gene (encodes a salicylate hydroxylase), and *nahH* gene (encodes a catechol 2, 3-dioxygenase). To construct the mutants that only possess the ability to perform one or a subset of metabolic steps in naphthalene degradation, we knocked out the four key genes one by one following a standard workflow, as shown in Figure S2. As a result, we totally obtained 16 mutants that possess the varied ability of naphthalene degradation. In the strain name, ‘1’ denotes that the related key gene is retained in the strain, thus it can carry out the corresponding step. In contrast, ‘0’ denotes that the related key gene is defective, thus it is not capable of carrying out the corresponding step. For example, strain *P. stutzeri* AN1000 (denoted as [1, 0, 0, 0] in the main Text) can convert naphthalene to 1,2-dihydroxynaphthalene, but cannot carry out the rest of the metabolic steps.

##### *S1.1.3 Genetic manipulation and strain validation*

The genetic manipulations were implemented by allele exchange using the suicide plasmid pK18mobsacB^9,10^. The constructed strains were validated by PCR and DNA sequencing. In addition, enzymic activity assays were performed to verify the phenotypes of different strains, following the methods reported before (Reference^11^ for naphthalene dioxygenase, Reference^12^ for 1, 2-dihydroxynaphthalene dioxygenase, Reference^13^ for salicylate 1-hydroxylase, and Reference^11^ for catechol 2, 3-dioxygenase). Moreover, we also mono-cultured these strains in the minimal medium supplemented with naphthalene, 1, 2- hydroxynaphthalene, salicylate, and catechol as the sole carbon source. As shown in Figure S1.3 and Figure S1.4, the genotypes of all the strains are consistent with our design.

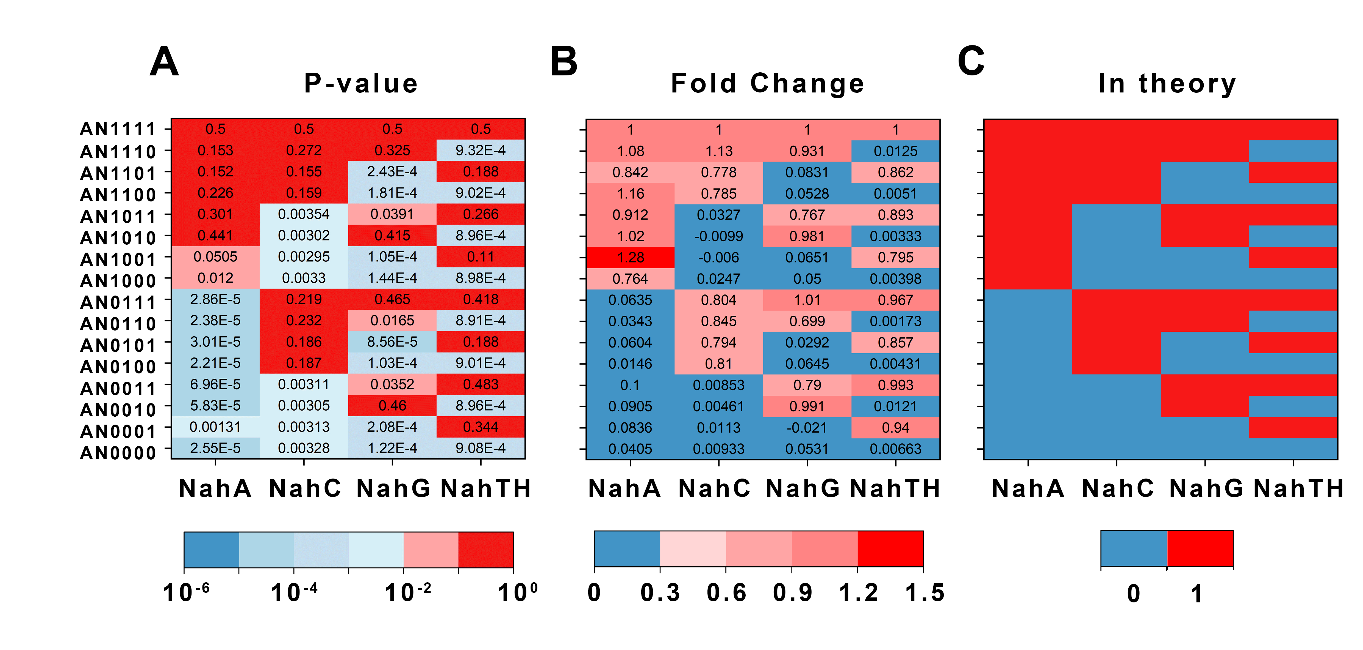

**Figure S1.3** Enzyme activity assays of all the strains used for constructing synthetic communities. (A) Two-tailed t-test showed a significant difference (*p*-value) in the enzyme activity of the four key enzymes between all other strains with *P. stutzeri* AN1111. (B) The average ratio of enzyme activity of four key enzymes in all strains to that of *P. stutzeri* AN1111. (C) Theoretical phenotypes of all strains. ‘0’ means the strain should not possess the enzyme activity, while ‘1’ means the strain should possess the enzyme activity.

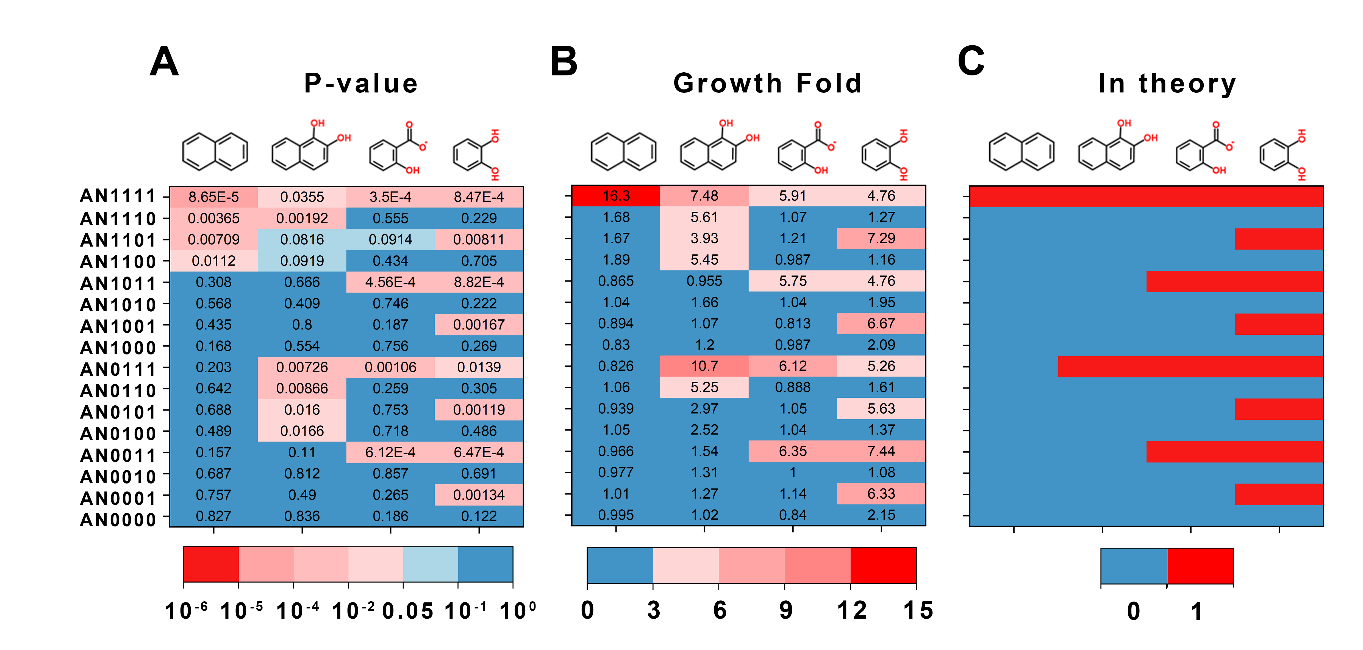

**Figure S1.4** The growth of all the strains using naphthalene and three intermediates as the sole carbon source. (A) Double-tailed T-test showed a significant difference (*P*-values) between the growth fold of all strains with that of *P. stutzeri* AN1111 in each carbon source. (B) relative growth fold between all strains with *P. stutzeri* AN1111 in each carbon source. (C) Theoretical phenotypes of all strains. ‘0’ means the strain should not grow in the related carbon source, while ‘1’ means the strain should grow in the related carbon source.

#### **1.2 Quantifying the relative frequencies of different strains in our synthetic consortia**

##### *S1.2.1 The principle of the measurement*

During the strain construction, a single base tag, A, T, G, or C, was introduced at each knockout site. These tags were summarized in Figure S1.5. If the gene was not knocked out, the corresponding locus was the original gene sequence. Therefore, one specific genotype possesses a unique combination of tags in its genomic regions of *nahA*, *nahC*, *nahG*, and *nahTH*. Accordingly, we could distinguish different genotypes from a mixed culture. After the co-culture of different genotypes in a consortium, we performed high-throughput amplicon sequencing of these four regions and calculated the relative frequencies of different tags in each region. Accordingly, the relative abundance of different genotypes can be solved using the following equations

*X*_[0, 1, 1, 1]_ + *X*_[0, 1, 0, 1]_ + *X*_[0, 0, 0, 1]_ + *X*_[0, 1, 1, 0]_ = *Y*_△_*_nahA::A_*	[S1]

*X*_[0, 0, 1, 1]_ + *X*_[0, 0, 1, 0]_ = *Y*_△_*_nahA::T_*	[S2]

*X*_[0, 1, 0, 0]_ = *Y*_△_*_nahA::C_*	[S3]

*X*_[0, 0, 0, 0]_ = *Y*_△_*_nahA::G_*	[S4]

*X*_[1, 1, 1, 0]_ + *X*_[1, 1, 0, 0]_ + *X*_[1, 0, 1, 1]_ + *X*_[1, 0, 1, 0]_ + *X*_[1, 0, 0, 0]_ + *X*_[1, 1, 0, 1]_ + *X*_[1, 0, 0, 1]_ + *X*_[1, 1, 1, 1]_ = *Y_nahA_*

[S5]

*Y_nahA::A_* + *Y_nahA::T_* + *Y_nahA::C_* + *Y_nahA::G_* + *Y_nahA_* = 100%	[S6]

*X*_[1, 0, 1, 1]_ + *X*_[0, 0, 1, 1]_ + *X*_[0, 0, 1, 0]_ + *X*_[1, 0, 1, 0]_ = *Y*_△_*_nahC::A_*	[S7]

*X*_[1, 0, 0, 1]_ = *Y*_△_*_nahC::T_*	[S8]

*X*_[0, 0, 0, 1]_ = *Y*_△_*_nahC::C_*	[S9]

*X*_[1, 0, 0, 0]_ + *X*_[0, 0, 0, 0]_ = *Y*_△_*_nahC::G_*	[S10]

*X*_[0, 1, 1, 1]_ + *X*_[1, 1, 0, 1]_ + *X*_[0, 1, 0, 1]_ + *X*_[0, 1, 1, 0]_ + *X*_[1, 1, 1, 0]_ + *X*_[1, 1, 0, 0]_ + *X*_[0, 1, 0, 0]_ + *X*_[1, 1, 1, 1]_ = *Y_nahC_*	[S11]

*Y*_△_*_nahC::A_* + *Y*_△_*_nahC::T_* + *Y*_△_*_nahC::C_* + *Y*_△_*_nahC::G_*+*Y_nahC_* = 100%	[S12]

*X*_[1, 1, 0, 1]_ + *X*_[1, 0, 0, 1]_ = *Y*_△_*_nahG::A_*	[S13]

*X*_[1, 1, 0, 0]_ + *X*_[0, 1, 0, 0]_ + *X*_[1, 0, 0, 0]_ + *X*_[0, 0, 0, 0]_ = *Y*_△_*_nahG::T_*	[S14]

*X*_[0, 1, 0, 1]_ + *X*_[0, 0, 0, 1]_ = *Y*_△_*_nahG::G_*	[S15]

*X*_[0, 1, 1, 1]_ + *X*_[1, 1, 1, 0]_ + *X*_[1, 0, 1, 1]_ + *X*_[0, 1, 1, 0]_ + *X*_[0, 0, 1, 1]_ + *X*_[1, 0, 1, 0]_ + *X*_[0, 0, 1, 0]_ + *X*_[1, 1, 1, 1]_ = *Y_nahG_*	[S16]

*Y* _△_ *_nahG::A_* + *Y* _△_ *_nahG::T_* + *Y* _△_ *_nahG::G_* + *Y_nahG_* = 100%	[S17]

*X*_[1, 1, 1, 0]_ + *X*_[1, 1, 0, 0]_ + *X*_[0, 1, 0, 0]_ + *X*_[1, 0, 0, 0]_ + *X*_[0, 0, 0, 0]_ = *Y*_△_*_nahTH::A_*	[S18]

*X*_[1, 0, 1, 0]_ = *Y*_△_*_nahTH::T_*	[S19]

*X*_[0, 0, 1, 0]_ = *Y*_△_*_nahTH::C_*	[S20]

*X*_[0, 1, 1, 0]_ = *Y*_△_*_nahTH::G_*	[S21]

*X*_[0, 1, 1, 1]_ + *X*_[1, 0, 1, 1]_ + *X*_[1, 1, 0, 1]_ + *X*_[0, 0, 1, 1]_ + *X*_[0, 1, 0, 1]_ + *X*_[1, 0, 0, 1]_ + *X*_[0, 0, 0, 1]_ + *X*_[1, 1, 1, 1]_ = *Y_nahTH_*	[S22]

*Y*_△_*_nahTH::A_* + *Y*_△_*_nahTH::T_* + *Y*_△_*_nahTH::C_* + *Y*_△_*_nahTH::G_* + *Y_nahTH_* = 100%	[S23]

Here, *Y* is the relative frequencies of different tags in a given genomic region. For example, *Y*_△_*_nahTH::A_*, *Y*_△_*_nahTH::T_*, *Y*_△_*_nahTH::C_*, and *Y*_△_*_nahTH::G_* represent the frequencies of the tag A, T, G, and C in the region of nahTH, respectively, while *Y_nahTH_* represents the relative frequency of the original sequence in this region. *X* represents the relative abundance of different genotypes in the consortium.

To obtain the values of different *Y*, high-throughput amplicon sequencing of these four regions was performed. For each region, two pairs of primers were designed and two PCR reactions were conducted, one for the amplification of the tags of the mutants and the other for the amplification of the original sequence. The size of amplified fragments was 400 bp. In addition, two control strains, one with a known tag mutant sequence and the other with a known tag wild type tag sequence, were added to the PCR system. The tag sequences of the control strain were 10 bases different from the original sequence, so they could be distinguished. After amplification, the two PCR products were combined for sequencing. Taking the nahTH gene region as an example, the values of *Y* could be calculated by the following equations

$\text{Y}_{\text{△}\text{nahTH::A}}\text{=}\frac{\text{(}\frac{\text{Z}_{\text{△}\text{nahTH::A}}}{\text{Z1}}\text{)}}{\left( \frac{\text{Z}_{\text{△}\text{nahTH::A}}}{\text{Z1}} \right)\text{+}\left( \frac{\text{Z}_{\text{△}\text{nahTH::T}}}{\text{Z1}} \right)\text{+}\left( \frac{\text{Z}_{\text{△}\text{nahTH::G}}}{\text{Z1}} \right)\text{+}\left( \frac{\text{Z}_{\text{△}\text{nahTH::C}}}{\text{Z1}} \right)\text{+(}\frac{\text{Z}_{\text{△}\text{nahTH}}}{\text{Z2}}\text{)}}$	[S24]

$\text{Y}_{\text{△}\text{nahTH::C}}\text{=}\frac{\text{(}\frac{\text{Z}_{\text{△}\text{nahTH::C}}}{\text{Z1}}\text{)}}{\left( \frac{\text{Z}_{\text{△}\text{nahTH::A}}}{\text{Z1}} \right)\text{+}\left( \frac{\text{Z}_{\text{△}\text{nahTH::T}}}{\text{Z1}} \right)\text{+}\left( \frac{\text{Z}_{\text{△}\text{nahTH::G}}}{\text{Z1}} \right)\text{+}\left( \frac{\text{Z}_{\text{△}\text{nahTH::C}}}{\text{Z1}} \right)\text{+(}\frac{\text{Z}_{\text{△}\text{nahTH}}}{\text{Z2}}\text{)}}$	[S25]

$\text{Y}_{\text{△}\text{nahTH::G}}\text{=}\frac{\text{(}\frac{\text{Z}_{\text{△}\text{nahTH::G}}}{\text{Z1}}\text{)}}{\left( \frac{\text{Z}_{\text{△}\text{nahTH::A}}}{\text{Z1}} \right)\text{+}\left( \frac{\text{Z}_{\text{△}\text{nahTH::T}}}{\text{Z1}} \right)\text{+}\left( \frac{\text{Z}_{\text{△}\text{nahTH::G}}}{\text{Z1}} \right)\text{+}\left( \frac{\text{Z}_{\text{△}\text{nahTH::C}}}{\text{Z1}} \right)\text{+(}\frac{\text{Z}_{\text{△}\text{nahTH}}}{\text{Z2}}\text{)}}$	[S26]

$\text{Y}_{\text{△}\text{nahTH}\text{∷}\text{C}}\text{=}\frac{\text{(}\frac{\text{Z}_{\text{△}\text{nahTH::C}}}{\text{Z1}}\text{)}}{\left( \frac{\text{Z}_{\text{△}\text{nahTH::A}}}{\text{Z1}} \right)\text{+}\left( \frac{\text{Z}_{\text{△}\text{nahTH::T}}}{\text{Z1}} \right)\text{+}\left( \frac{\text{Z}_{\text{△}\text{nahTH::G}}}{\text{Z1}} \right)\text{+}\left( \frac{\text{Z}_{\text{△}\text{nahTH::C}}}{\text{Z1}} \right)\text{+(}\frac{\text{Z}_{\text{△}\text{nahTH}}}{\text{Z2}}\text{)}}$	[S27]

$\text{Y}_{\text{△}\text{nahTH}}\text{=}\frac{\text{(}\frac{\text{Z}_{\text{△}\text{nahTH}}}{\text{Z2}}\text{)}}{\left( \frac{\text{Z}_{\text{△}\text{nahTH::A}}}{\text{Z1}} \right)\text{+}\left( \frac{\text{Z}_{\text{△}\text{nahTH::T}}}{\text{Z1}} \right)\text{+}\left( \frac{\text{Z}_{\text{△}\text{nahTH::G}}}{\text{Z1}} \right)\text{+}\left( \frac{\text{Z}_{\text{△}\text{nahTH::C}}}{\text{Z1}} \right)\text{+(}\frac{\text{Z}_{\text{△}\text{nahTH}}}{\text{Z2}}\text{)}}$	[S28]

Here, *Z_nahTH::A_*, *Z_nahTH::T_*, *Z_nahTH::C_*, and Z*_nahTH::G_* represent the number of reads containing the tags A, T, G, and C, respectively and *Z1* is the number of reads of the mutant control strain. *Z_nahTH_* is the number of reads containing wild-type sequences, and *Z2* is the frequency of the number of reads of the wild-type control strain. This workflow was summarized in Figure S1.6.

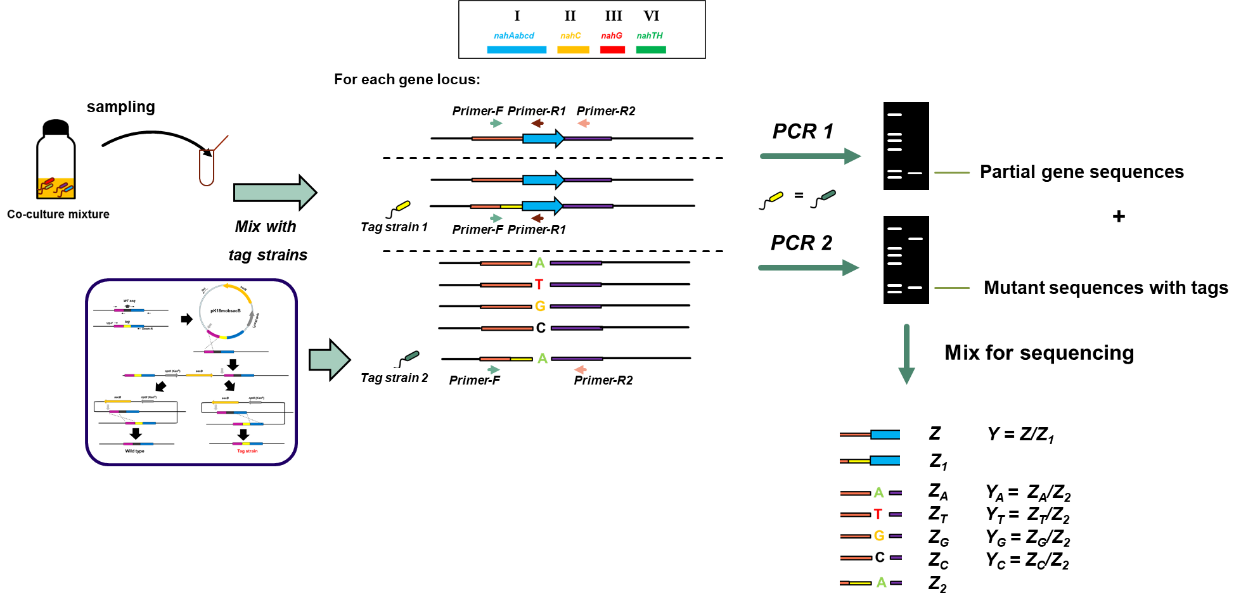

**Figure S1.6** Preparation of samples for high-throughput sequencing

##### *S1.2.2 Experimental protocols and data analysis*

The PCR reactions were performed using 2 μL of the culture of the consortia (OD_600_: 0.25) plus 1 μL of the suspension of the two control strains (OD_600_: 0.25; 1 to 1 ratio mixture) as the templates. 2X Taq Plus Master Mix II (Vazyme, Nanjing, China) was used for amplification following the standard protocol. The DNA in the derived PCR products was quantified and mixed equally. PE250 mode of Illumina HiSeq 2500 high-throughput sequencing platform was adopted for sequencing. During the sequencing, 96 PCR samples were mixed into a library, and a total of 48 libraries were established. Approximately 20 million reads were obtained. Therefore, around 7,000 reads could be allocated to each sample on average for subsequent analysis (Figure 1.7). The data were analyzed using the Qiime2 platform (version 2019.4) following the standard workflow to obtain the values of *Z*. Custom Mathematica scripts were further used the solve the above-mentioned equations to obtain the relative frequencies of different genotypes in our consortia.

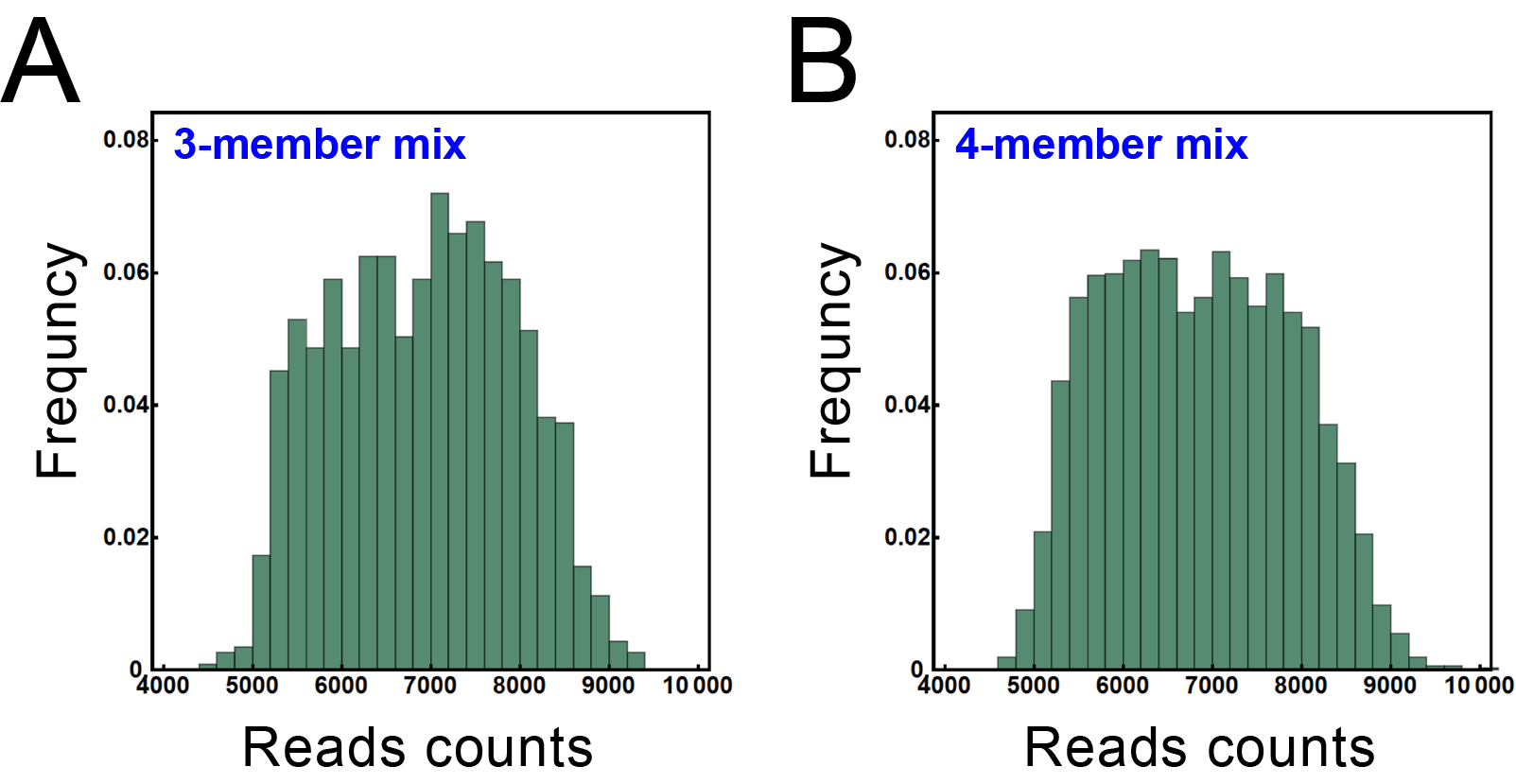

**Figure S1.7** Distribution of the reads number of the samples for three-member consortia (A) and four-member consortia (B).

##### *S1.2.3 Test of the Methodology*

To test whether our method is robust to quantify the relative frequencies of different strains in our synthetic consortia, we selected several typical two-member, three-member, four-member, 14-member, 15-member, and 16-member consortia and mixed their members with a series of proportions. We performed the measurements of the relative frequencies following the workflow proposed previously to test whether the frequencies obtained by sequencing were consistent with that of what we premixed. The results showed a high consistency between the measured frequencies and the expected frequencies. In the case of two-, three-, and four-member consortia, the linear correlation coefficients between measured frequencies and the expected frequencies are all above 0.9 (Figure S1.8A-C). With the increase of strain diversity, the inconsistency slightly increased. However, the sequencing method still showed good robustness even with the combination of 14, 15, and 16 members (Figure S1.8D). This result indicated that our method is robust to quantify the relative frequencies of different strains in our synthetic consortia.

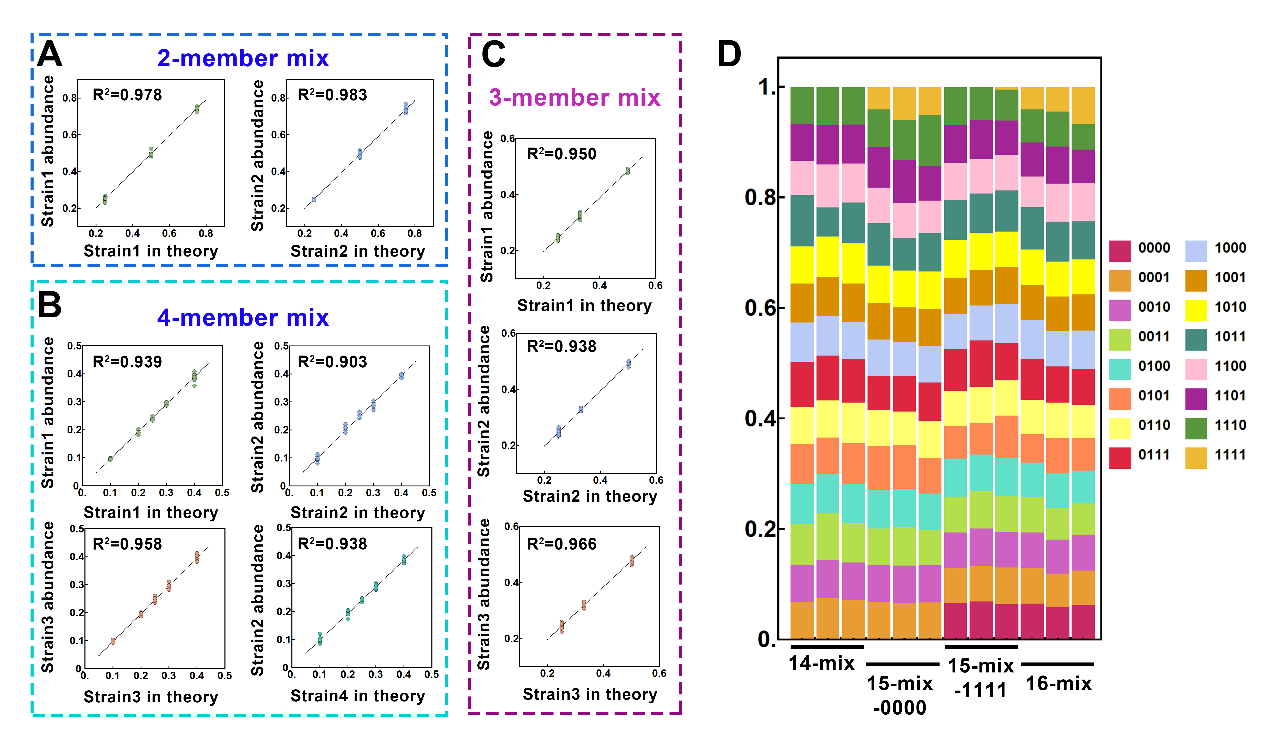

**Figure S1.8** Feasibility of the high-throughput sequencing method. (A) Two-member combinations, 0001 and 1110,0010, 1101,0100 and 1101,1000 and 0111, were selected and mixed with three ratios, 0.75-0.25, 0.50-0.50 and 0.25.0.75, and sequenced with two repeated mix samples. The figure shows the correlation between theoretical frequencies and measured frequencies of strain 1 and strain 2 in all combinations. (B) Three-member combinations, (0001, 0010, 1100), (1000, 0100 and 0011), and (1110, 1011, 1101) were selected and mixed with four ratios, 0.25-0.25-0.50, 0.25-0.50-0.25, 0.50-0.25-0.25, and 0.33-0.33-0.33 for each combination, and sequenced with two repeated mix samples. The figure shows the correlation between theoretical abundance and measured abundance of strain 1, strain 2, and strain 3 in all combinations. (C) Four-member combinations, (0001, 0010, 0100, and 1000), (1110, 1101, 1011, and 0111), and (1100, 0110, 0011, and 1001), were selected and mixed with six ratios 0.10-0.20-0.30-0.40, 0.20-0.30-0.40-0.10, 0.30-0.40-0.10-0.20, 0.40-0.10-0.20-0.30, 0.40-0.10-0.40-0.10, 0.25-0.25-0.25.0-25 and sequenced with two repeated mix samples. The figure shows the correlation between theoretical abundance and measured abundance of strain 1, strain 2, strain 3, and strain 4 in all combinations. (D) Based on the seed bank of 16 strains, 14 strains (except 0000 and 1111) were mixed in equal proportion, two 15 strains were mixed in equal proportion (except 0000 or 1111), and 16 strains were mixed in equal proportion. Each case was sequenced three times. The measured sequencing results of these combinations are shown in the figure.

#### **1.3 Definition and measurements of the four quantitative indexes**

##### *S1.3.1 Functional redundancy level*

The functional redundancy level (FR) of one consortium was defined following one previous work^14^. FR is defined based on functional dissimilarities among species, which is quantified using a formula:

$\text{FR}\text{ }\text{=}\text{ }\frac{\text{D}\text{ }\text{-}\text{ }\text{Q}}{\text{D}}\text{ }$	[S29]

Here, *D* represents the Simpson index^15^, given by

$\text{D }\text{= 1 - }\sum_{\text{i}} \text{p}_{\text{i}}\text{(1}\text{ }\text{-}{\text{ }\text{p}}_{\text{i}}\text{)}\text{ }\text{ }$	[S30]

where $\text{p}_{\text{i}}$ is the initial frequency of the *ith* strain residing in the consortium (that is, $\text{p}_{\text{i}}\text{ }\text{=}\text{ }\text{0.25}$ for a four-member consortium). *Q* represents the average dissimilarity of an individual from the whole consortium, which is a measure of the functional dissimilarity of the consortium. *Q* is given by^14^

$\text{Q}\text{ }\text{= 1 - }\sum_{\text{i}} \text{p}_{\text{i}}\sum_{\text{j}} \text{p}_{\text{j}}\text{δ}_{\text{i,j}}\text{ }\text{ }$	[S31]

$\text{δ}_{\text{i,j}}$ is the pairwise functional dissimilarity between strains *i* and *j* with a range of 0 ~ 1. Bray–Curtis dissimilarities were used to estimate $\text{δ}_{\text{i,j}}$. For any given pair of strains with their genotype characterized by bit strings, the Bray–Curtis dissimilarities of the bit string pairs were calculated to estimate $\text{δ}_{\text{i,j}}$.

##### *S1.3.2 The average metabolic burden*

*The average metabolic burden (AMB)* of one consortium represents the average value of metabolic burdens of the strains involved in the consortium. The metabolic burden of one mutant strain is defined as the relative fitness between the autonomous strain [1, 1, 1, 1] and the corresponding strain, namely *F_g_*, in which *g* represents the genotype of the strain. Following this definition, the metabolic burden of [1, 1, 1, 1], *F*_[1, 1, 1, 1]_, equals 1.0, representing the maximum metabolic burden of all the strains. The metabolic burdens of other strains were then determined by performing pairwise competitive fitness assays in the minimal medium supplemented with pyruvate (0.5 % w/v; the end product of naphthalene degradation; Both strains can grow on) as the sole carbon source and 1 mM IPTG. Note that the expression of naphthalene degradation genes was induced by IPTG in our strains, so these genes were expressed in this culture condition even without the addition of naphthalene, imposing a metabolic burden on the strains. Strain [1, 1, 1, 1] was labeled with eGfp and the other strain in the system was labeled with mBeRFP. The relative frequencies of the two strains in this co-culture were measured by CFU counting. The calculation of relative fitness follows previous studies^16,17^

$\text{F}\text{g}\text{ = }\frac{\ln\frac{\text{N}_{\text{[1, 1, 1, 1]}\text{,}\text{ t}}}{\text{N}_{\text{[1, 1, 1, 1]}\text{,}\text{ 0}}}}{\ln\frac{\text{N}_{\text{g,t}}}{\text{N}_{\text{g,0}}}}\text{ }\text{ }$	[S32]

Here, $\text{N}_{\left[ \text{1, 1, 1, 1} \right]\text{, t}}$ and $\text{N}_{\left[ \text{1, 1, 1, 1} \right]\text{, 0}}$represent the CFU number of strain [1, 1, 1, 1] at the early stationary phase (18 h) and the beginning (0 h); similarly, $\text{N}_{\text{g,t}}$ and $\text{N}_{\text{g,0}}$represent the CFU number of focal strain at the early stationary phase and the beginning. $\text{F}\text{g}$ values of different strains are summarized in Table S1.1. AMB of one consortium was estimated by the average number of metabolic burdens of the strains involved in the consortium.

Table S1.1 $\text{F}\text{g}$ values of all strains

| Strains | $\text{F}\text{g}$ |
| --- | --- |
| *P. stutzeri* AN1111 | 1.000 |
| *P. stutzeri* AN0000 | 0.819 |
| *P. stutzeri* AN0001 | 0.869 |
| *P. stutzeri* AN0010 | 0.862 |
| *P. stutzeri* AN0011 | 0.897 |
| *P. stutzeri* AN0100 | 0.844 |
| *P. stutzeri* AN0101 | 0.862 |
| *P. stutzeri* AN0110 | 0.862 |
| *P. stutzeri* AN0111 | 0.897 |
| *P. stutzeri* AN1000 | 0.925 |
| *P. stutzeri* AN1001 | 0.868 |
| *P. stutzeri* AN1010 | 0.944 |
| *P. stutzeri* AN1011 | 0.980 |
| *P. stutzeri* AN1100 | 0.913 |
| *P. stutzeri* AN1101 | 0.948 |
| *P. stutzeri* AN1110 | 0.961 |

##### *S1.3.3 The average functional capacity*

The average functional capacity (AFC) of one consortium represents the average value of the functional capacity of the strains involved in the consortium. The functional capacity of one strain indicates how many metabolic steps this strain can perform, which is simply quantified by counting the number of “1” in the bit strings vector of the strain. The AFC was then estimated by the average value of functional capacity of the strains involved in the consortium.

##### *S1.3.4 The transport capacity of metabolites*

The transport capacity of metabolites (TCM) of one consortium represents the overall capacity of the transport of all intermediate metabolites of one consortium. Three intermediate metabolites are involved in our experimental design, in which the naphthalene degradation pathway was partitioned into four steps. First, for a strain that can perform the step to produce an intermediate, as well as the step to further convert the intermediate, the transport of the intermediate occurred intracellularly, which is rapid. Therefore, a value of 1 was assigned to assess the capacity of this strain to transport this intermediate. Second, for a strain that can perform the step to produce an intermediate but cannot further convert the intermediate, the intermediate must transport outside of the cell and then uptake by other strains that can convert this intermediate to continue the pathway. This transport process is slower, so a value of 0.5 was assigned to assess the transport capacity. Third, for a strain that cannot perform the step to produce an intermediate, we assign a value of 0, which means that the strain has no contribution to the production and transport of the intermediate. The overall *TCM* of an intermediate of one consortium was then defined as the average capacity to transport the intermediate of the strains involved in the consortium. Finally, The *TCM* of one consortium was evaluated as the average value of the *TCM* of all three intermediates.

#### **1.4 Derivation of the mathematical model**

##### *S1.4.1 The assumptions of the model*

Our mathematical model uses ordinary differential equations (ODEs) to characterize the dynamics of four-member MDOL consortia possessing functional redundancy (MCFRs). The definitions and dimensionless methods of all the variables and parameters used in the ODEs are listed in Table S1-S2. Same to our experimental system (Figure 1 and Figure S1), we assume that a degradation pathway is divided into four steps and can be collectively performed by four populations. Each population can perform one, two, or three metabolic step(s) of the pathway. One initial substrate (S), three intermediates (I1, I2, I3), and an end product (P) are included. For simplicity, the model was built based on seven simple assumptions identical to our previous study^18^. We restated these assumptions as follows

(1) The systems for both configurations are well mixed in each compartment (inside a cell or in the extracellular space).

(2) Transport of the metabolites, including substrate (S), intermediates (I), and end product (P), across the cell membrane, occurs via passive diffusion at a rate proportional to the concentration gradient between two compartments.

(3) Metabolic reactions were performed intracellularly.

(4) The spontaneous degradation of I and P is significantly slower than kinetic reactions and thus its rate may be set to approximately 0.

(5) The supply of S in the system is sufficient, and thus the rate of S consumption is independent of the concentration of S.

(6) The intracellular accumulation of I and P in the system is negligible, and thus the rates of the two metabolic reactions follow the first-order kinetics.

(7) The end product of the pathway is the sole carbon source that all populations use for growth.

In our previous study^18^, we have shown that the model built based on these assumptions can accurately predict the dynamics of our synthetic consortia performing MDOL to degrade naphthalene. In that study, we also performed detailed analyses to address how relaxing each assumption can affect the results of the simulations.

In this study, we modified that model by adding the assumption of functional redundancy. This assumption is formulated by conceptualizing the genotypes of the strains using bit strings containing “0” and “1” ($\text{ϵ}_{\text{k}}$).

$\text{ϵ}_{\text{k}}\text{=}\left[ \text{ϵ}_{\text{k,1}}\text{,}\text{ }\text{ϵ}_{\text{k,2}}\text{,}\text{ }\text{ϵ}_{\text{k,3}}\text{,}\text{ }\text{ϵ}_{\text{k,4}} \right]$	$\text{ }\text{ }$	[S33]

in which

$\text{ϵ}_{\text{k,j}}\text{=}\text{1} \text{or} \text{0}$	$\text{ }\text{ }$	[S34]

Here, 1 denotes that the corresponding population is capable of executing the *j*th step, and 0 denotes it cannot.

##### *1.4.2 Equations regarding Intermediate and product dynamics*

The ODEs describing the dynamics of Intermediates and product concentrations are given by

$\frac{\text{d}\text{S}_{\text{,k,in}}}{\text{dt}}\text{∙}\text{V}_{\text{c}}\text{ }\text{=}\text{ }\text{-}\text{ϵ}_{\text{k,1}}\frac{\text{k}_{\text{1}}\text{E}_{\text{1}}}{\text{K}_{\text{1}}\text{+}\text{S}_{\text{1,}\text{k,}\text{in}}}\text{S}_{\text{k,in}}\text{ }\text{+}\text{ }\text{r}_{\text{1}}\text{∙}\text{V}_{\text{c}}\text{∙}\text{ }\left( \text{S}_{\text{out}} \text{-}\text{ }\text{S}_{\text{k,in}} \right)$	[S35]

$\frac{\text{d}\text{I}_{\text{1,k,in}}}{\text{dt}}\text{∙}\text{V}_{\text{c}}\text{ = }\text{ϵ}_{\text{k,1}}\frac{\text{k}_{\text{1}}\text{E}_{\text{1}}}{\text{K}_{\text{1}}\text{+}\text{S}_{\text{1,}\text{k,}\text{in}}}\text{S}_{\text{k,in}} \text{- }\text{ϵ}_{\text{k,2}}\frac{\text{k}_{\text{2}}\text{E}_{\text{2}}}{\text{K}_{\text{2}}\text{+}\text{I}_{\text{2,k,in}}}\text{I}_{\text{1,}\text{k,}\text{in}}\text{ + }\text{r}_{\text{2}}\text{∙}\text{V}_{\text{c}}\text{∙ }\left( \text{I}_{\text{1,out}}\text{ }\text{- }\text{I}_{\text{1,k,in}} \right)$	[S36]

$\frac{\text{d}\text{I}_{\text{2,k,in}}}{\text{dt}}\text{∙}\text{V}_{\text{c}}\text{ }\text{=}\text{ }\text{ϵ}_{\text{k,2}}\frac{\text{k}_{\text{2}}\text{E}_{\text{2}}}{\text{K}_{\text{2}}\text{+}\text{I}_{\text{2,}\text{k,}\text{in}}}\text{I}_{\text{1,in}} \text{-}\text{ }\text{ϵ}_{\text{k,3}}\frac{\text{k}_{\text{3}}\text{E}_{\text{3}}}{\text{K}_{\text{3}}\text{+}\text{I}_{\text{3,}\text{k,}\text{in}}}\text{I}_{\text{2,}\text{k,}\text{in}}\text{ }\text{+}\text{ }\text{r}_{\text{3}}\text{∙}\text{V}_{\text{c}}\text{∙}\text{ }\left( \text{I}_{\text{2,out}} \text{-}\text{ }\text{I}_{\text{2,k,in}} \right)$	[S37]

$\frac{\text{d}\text{I}_{\text{3,k,in}}}{\text{dt}}\text{∙}\text{V}_{\text{c}}\text{ }\text{=}\text{ }\text{ϵ}_{\text{k,3}}\frac{\text{k}_{\text{3}}\text{E}_{\text{3}}}{\text{K}_{\text{3}}\text{+}\text{I}_{\text{3,}\text{k,}\text{in}}}\text{I}_{\text{2,in}} \text{-}\text{ }\text{ϵ}_{\text{k,4}}\frac{\text{k}_{\text{4}}\text{E}_{\text{4}}}{\text{K}_{\text{4}}\text{+}\text{I}_{\text{4,}\text{k,}\text{in}}}\text{I}_{\text{3,}\text{k,}\text{in}}\text{ }\text{+}\text{ }\text{r}_{\text{4}}\text{∙}\text{V}_{\text{c}}\text{∙}\text{ }\left( \text{I}_{\text{3,out}} \text{-}\text{ }\text{I}_{\text{3,k,in}} \right)$	[S38]

$\frac{\text{d}\text{p}_{\text{k,in}}}{\text{dτ}}\text{∙}\text{V}_{\text{c}}\text{ }\text{=}\text{ }\text{ϵ}_{\text{k,4}}\frac{\text{k}_{\text{4}}\text{E}_{\text{4}}}{\text{K}_{\text{4}}\text{+}\text{I}_{\text{4,}\text{k,}\text{in}}}\text{I}_{\text{3,in}} \text{-}\text{ }\frac{\text{k}\text{g}}{\text{K}_{\text{g}}\text{+}\text{P}_{\text{k,in}}}\text{∙}\text{P}_{\text{k,in}}\text{ }\text{+}\text{ }\text{ }\text{r}_{\text{5}}\text{∙}\text{V}_{\text{c}}\text{∙}\left( \text{P}_{\text{out}} \text{-}\text{ }\text{P}_{\text{k,in}} \right)$	[S39]

$\frac{\text{d}\text{S}_{\text{out}}}{\text{dt}}\text{∙}\text{V}_{\text{c}}\text{ }\text{=}\text{ }\text{-}\sum_{\text{k=1}}^{\text{4}} \text{X}_{\text{k}}\text{∙}\text{r}_{\text{1}}\text{∙}\left( \text{S}_{\text{out}} \text{-}\text{ }\text{S}_{\text{k,in}} \right)$	[S40]

$\frac{\text{d}\text{I}_{\text{j,out}}}{\text{dt}}\text{∙}\text{V}_{\text{c}}\text{ }\text{=}\text{ }\text{-}\sum_{\text{k=1}}^{\text{4}} \text{X}_{\text{k}}\text{∙}\text{r}_{\text{j}\text{+1}}\text{∙}\left( \text{I}_{\text{j,out}}\text{ }\text{- }\text{I}_{\text{j,k,in}} \right)$	[S41]

$\frac{\text{d}\text{P}_{\text{out}}}{\text{dt}}\text{∙}\text{V}_{\text{c}}\text{=-}\sum_{\text{k=1}}^{\text{4}} \text{X}_{\text{k}}\text{∙}\text{γ}_{\text{5}}\text{∙}\left( \text{P}_{\text{out}} \text{-}\text{ }\text{P}_{\text{k,in}} \right)$	[S42]

*j* = 1 ~ 3 (meaning three intermediates) and *k* = 1 ~ 4 (meaning four genotypes involved in the consortium). Therefore, the system contains 25 equations. According to assumptions (4) and (5), We further assumed that $\text{S}_{\text{k,in}}\text{ ≫}\text{ }\text{K}_{\text{1}}$,$\text{K}_{\text{j+1}}\text{ ≫}\text{I}_{\text{j,}\text{k, }\text{in}}$, $\text{K}_{\text{g}}\text{ ≫}\text{P}_{\text{k,in}}$, which simplified the model to a linear system,

$\frac{\text{d}\text{S}_{\text{,k,in}}}{\text{dt}}\text{ }\text{=}\text{ }\text{-}\text{ϵ}_{\text{k,1}}\text{k}_{\text{1}}\text{E}_{\text{1}}\text{ }\text{+}\text{ }\text{r}_{\text{1}}\text{∙}\text{ }\left( \text{S}_{\text{out}} \text{-}\text{ }\text{S}_{\text{k,in}} \right)$	[S43]

$\frac{\text{d}\text{I}_{\text{1,k,in}}}{\text{dt}}\text{ }\text{=}\text{ }\text{ϵ}_{\text{k,1}}\text{k}_{\text{1}}\text{E}_{\text{1}} \text{-}\text{ }\text{ϵ}_{\text{k,2}}\frac{\text{k}_{\text{2}}\text{E}_{\text{2}}}{\text{K}_{\text{2}}}\text{I}_{\text{1,}\text{k,}\text{in}}\text{ }\text{+}\text{ }\text{r}_{\text{2}}\text{∙}\text{ }\left( \text{I}_{\text{1,out}} \text{-}\text{ }\text{I}_{\text{1,k,in}} \right)$	[S44]

$\frac{\text{d}\text{I}_{\text{2,k,in}}}{\text{dt}}\text{ }\text{=}\text{ }\text{ϵ}_{\text{k,2}}\frac{\text{k}_{\text{2}}\text{E}_{\text{2}}}{\text{K}_{\text{2}}}\text{I}_{\text{1,in}} \text{-}\text{ }\text{ϵ}_{\text{k,3}}\frac{\text{k}_{\text{3}}\text{E}_{\text{3}}}{\text{K}_{\text{3}}}\text{I}_{\text{2,}\text{k,}\text{in}}\text{ }\text{+}\text{ }\text{r}_{\text{3}}\text{∙}\text{ }\left( \text{I}_{\text{2,out}} \text{-}\text{ }\text{I}_{\text{2,k,in}} \right)$	[S45]

$\frac{\text{d}\text{I}_{\text{3,k,in}}}{\text{dt}}\text{ }\text{=}\text{ }\text{ϵ}_{\text{k,3}}\frac{\text{k}_{\text{3}}\text{E}_{\text{3}}}{\text{K}_{\text{3}}}\text{I}_{\text{2,in}} \text{-}\text{ }\text{ϵ}_{\text{k,4}}\frac{\text{k}_{\text{4}}\text{E}_{\text{4}}}{\text{K}_{\text{4}}}\text{I}_{\text{3,}\text{k,}\text{in}}\text{ }\text{+}\text{ }\text{r}_{\text{4}}\text{∙}\text{ }\left( \text{I}_{\text{3,out}} \text{-}\text{ }\text{I}_{\text{3,k,in}} \right)$	[S46]

$\frac{\text{d}\text{p}_{\text{k,in}}}{\text{dτ}}\text{∙}\text{ }\text{=}\text{ }\text{ϵ}_{\text{k,4}}\frac{\text{k}_{\text{4}}\text{E}_{\text{4}}}{\text{K}_{\text{4}}}\text{I}_{\text{3,in}} \text{-}\text{ }\frac{\text{k}\text{g}}{\text{K}_{\text{g}}}\text{∙}\text{P}_{\text{k,in}}\text{ }\text{+}\text{ }\text{ }\text{r}_{\text{5}}\text{∙}\left( \text{P}_{\text{out}} \text{-}\text{ }\text{P}_{\text{k,in}} \right)$	[S47]

$\frac{\text{d}\text{S}_{\text{out}}}{\text{dt}}\text{∙}\text{V}_{\text{c}}\text{ }\text{=}\text{ }\text{-}\sum_{\text{k=1}}^{\text{4}} \text{X}_{\text{k}}\text{∙}\text{r}_{\text{1}}\text{∙(}\text{S}_{\text{out}\text{ }}\text{-}\text{ }\text{S}_{\text{k,in}}\text{)}$	[S48]

$\frac{\text{d}\text{I}_{\text{j,out}}}{\text{dt}}\text{∙}\text{V}_{\text{c}}\text{ }\text{=}\text{ }\text{-}\sum_{\text{k=1}}^{\text{4}} \text{X}_{\text{k}}\text{∙}\text{r}_{\text{j}\text{+1}}\text{∙(}\text{I}_{\text{j,out}} \text{-}\text{ }\text{I}_{\text{j,k,in}}\text{)}$	[S49]

$\frac{\text{d}\text{P}_{\text{out}}}{\text{dt}}\text{∙}\text{V}_{\text{c}}\text{=-}\sum_{\text{k=1}}^{\text{4}} \text{X}_{\text{k}}\text{∙}\text{γ}_{\text{5}}\text{∙(}\text{P}_{\text{out}} \text{-}\text{ }\text{P}_{\text{k,in}}\text{)}$	[S50]

The equations were then non-dimensionalized following the method described in Table S1 and Table S2, which formalized a simple, non-dimensionalized model, presented in the Method section of the main text as equations [2] - [9].

##### *S1.4.3 Equations regarding population growth dynamics*

The ODEs describing the growth of the *k*th population are given by

$\frac{\text{d}\text{X}_{\text{k}}}{\text{dt}}\text{=}\text{g}_{\text{k}}\text{X}_{\text{k}}\left( \text{1-}\frac{\sum_{\text{k=1}}^{\text{4}} \text{X}_{\text{k}}}{\text{N}_{\text{m}}} \right)$	[S51]

Since all populations can only use the end product P as the sole carbon source (assumption (7)), the growth rate of the *k*th population ($\text{g}_{\text{k}}$), are calculated by

$\text{g}_{\text{k}}\text{=}\frac{\text{k}\text{g}\text{P}_{\text{k,in}}}{\text{K}_{\text{g}}\text{+}\text{P}_{\text{k,in}}}\text{Y}\text{M}_{\text{k}}$	[S52]

Similarly, equation [S52] was simplified following the assumption (4), that is $\text{K}_{\text{g}}\text{ ≫}\text{P}_{\text{k,in}}$, and thus simplified as

$\text{g}_{\text{k}}\text{=}\frac{\text{k}\text{g}\text{P}_{\text{k,in}}}{\text{K}_{\text{g}}}\text{Y}\text{M}_{\text{k}}$	[S53]

In particular, the metabolic burden of the kth population, $\text{M}_{\text{k}}$, was modeled following a previous study^19^, given by

$\text{M}_{\text{k}}\text{=}\frac{\text{K}_{\text{burd}}^{\text{h}}}{\text{K}_{\text{burd}}^{\text{h}}\text{+}\left( \sum_{\text{j=1}}^{\text{4}} \text{ϵ}_{\text{k,j}}\text{E}_{\text{j}} \right)^{\text{h}}}$	[S54]

The equation [53] was non-dimensionalized following the method described in Table S1 and Table S2, which formalized the non-dimensionalized equation presented in the Method section of the main text as equations [10]. In addition, the equation [54] was non-dimensionalized following the method described in Table S1 and Table S2, obtaining an equation defining the non-dimensionalized metabolic burden, $\text{m}_{\text{k}}$, given by

$\text{m}_{\text{k}}\text{=}\frac{\text{1}}{\text{1}\text{ }\text{+}\left( \sum_{\text{j=1}}^{\text{4}} \text{ϵ}_{\text{k,j}}\text{c}_{\text{j}}\text{e}_{\text{j}} \right)^{\text{h}}}$	[S55]

in which

$\text{c}_{\text{j}}=\frac{\text{K}_{\text{j}}}{\text{K}_{\text{burd}}}$	[S56]

##### *S1.4.4 Simulation protocols*

Numerical simulations of the model were performed using the NDsolve function of Wolfram Mathematica (version 12.0). The numerical solutions of all the variables, including the dynamics of mass (S, I, P) concentration and biomass, were recorded for further analyses. To measure the average growth rates of each modeled consortium, the dynamics of total biomass ($\sum_{\text{k=1}}^{\text{4}} \text{X}_{\text{k}}$, reflecting the growth curve of the consortium) were fitted to Logistic Function^20^. The degradation rate of S of each modeled consortium was calculated at the time point when the growth of the consortium just reached the stationary phase in our simulations. These measurements are consistent with those in our experiments.

To mathematically predict the experimental results, parameter values matching our experimental system (Table S1) were applied to the model. The dynamics of all the 860 MCFRs, one rigorous MDOL consortium, and one autonomous population (meaning different combinations of $\text{ϵ}_{\text{k}}$) were simulated. The average growth rates and degradation rates of all these systems were calculated, normalized, and compared to the experimental results.

To generalize the experimental findings in more pathway conditions, 138500 parameter sets were generated. In total, the model contains 24 parameters that may affect the simulation results. We parameterized eight of them, namely $\text{cp}$, $\text{y}$, $\text{ρ}$, *s_0_*, and *x_k,0_* (*k* = 1 ~ 4), using the values experimentally measured or the same as our experimental set-up. The rest of the 16 parameters, namely $\text{α}_{\text{i}}$ (*i* = 2 ~ 4), $e_{\text{i}}$ (*i* = 1 ~ 4), $c_{\text{j}}$ (*j* = 1 ~ 4), and $\text{γ}$*_j_* (*j* = 1 – 5), were parameterized by randomly picking the values from the given ranges obtained from literature search or experimental measurements (Table S1). The rationale behind this set-up is that these 16 parameters directly affect the metabolic burden of strains, the functional capacity of the consortium, and the transport capacity of metabolites in the consortium, which are the important factors given by our hypothesis and tested in the experiments. We thus focus on the effects of those important parameters on the functions of the consortia. In total, we generated 138500 random parameter sets. We adopted each parameter set to model the dynamics of all the 862 MCFRs, one rigorous MDOL consortium, and one autonomous population. The average growth rates and degradation rates of different systems were calculated and compared to explore: (1) whether there are any MCFRs show better function than the rigorous MDOL consortium and the autonomous population; (2) the set-up of the MCFRs with the best function; (3) the effects of the main parameters on the functions of the MCFRs.

All the simulations, as well as the downstream analysis, were performed using custom Mathematica scripts. The source codes used for the model analysis are publicly available (<https://github.com/WMXgg/MDOLcode/tree/master/MDOL-redun>).

### S2 Supplementary Tables and Figures

**Table S1** Definitions, dimensionless methods, and values of the parameters involved in our model.

| **Parameter** | | **Definitions** | **Values for prediction** | **Source** | **ND parameter** | **ND value for prediction** | **The range for parameter set generation** |
| --- | --- | --- | --- | --- | --- | --- | --- |
| $\text{K}_{\text{j}}$ | Michaelis-Menten constant for the *jth* reaction (*j* = 1 ~ 4). | | $\text{K}_{\text{1}}$ = 9.2×10^-7^ M  $\text{K}_{\text{2}}$ = 3.4×10^-5^ M  $\text{K}_{\text{3}}$ = 9.6×10^-6^ M  $\text{K}_{\text{4}}$ = 1.5×10^-6^ M | ^21^ | $\text{e}_{\text{j}}\text{=}\text{E}_{\text{j}}\text{/}{\text{(}\text{K}}_{\text{j}}\text{)}$  $\text{α}_{\text{j}}\text{=}\text{k}_{\text{j}}\text{/}\text{k}_{\text{1}}$*,* $\text{α}_{\text{1}}\text{=}\text{1}$ | $\text{e}_{\text{1}}$ = 2.2  $\text{e}_{\text{2}}$ = 0.059  $\text{e}_{\text{3}}$ = 0.21  $\text{e}_{\text{4}}$ = 1.3  $\text{α}_{\text{1}}$ = 1  $\text{α}_{\text{2}}$ = 12.0  $\text{α}_{\text{3}}$ = 2.5  $\text{α}_{\text{4}}$ = 110 | $\text{e}_{\text{j}}\text{ ϵ (0, 10]}$  $\text{α}_{\text{j}}\text{ ϵ }\text{[}\text{10}\text{-3}\text{, }\text{10}\text{3}\text{]}$ |
| $\text{E}_{\text{j}}$ | Concentration of the *jth* enzyme (*j* =1 ~ 4). | | 2×10^-6^ mol⸱L^-1^ | ^21-23^ |  |  |  |
| $\text{k}_{\text{j}}$ | Specific rate of the *jth* reaction (*j* =1 ~ 4). | | $\text{k}_{\text{1}}$ = 1.82 s^-1^  $\text{k}_{\text{2}}$ = 22.7 s^-1^  $\text{k}_{\text{3}}$ = 4.6 s^-1^  $\text{k}_{\text{4}}$ = 200 s^-1^ | ^24^ |  |  |  |
| $\text{r}_{\text{j}}$ | Diffusivity of the substrate, three intermediates, and final product (*j* =1 ~ 5). | | $\text{r}_{\text{1}}$ = 3.35 s^-1^  $\text{r}_{\text{2}}$ = 3020 s^-1^  $\text{r}_{\text{3}}$ = 26.3 s^-1^  $\text{r}_{\text{4}}$ = 4.27 s^-1^  $\text{r}_{\text{5}}$ = 25.1 s^-1^ | ^25^ | $\text{γ}_{\text{j}}\text{=}\text{r}_{\text{j}}\text{/}\text{k}_{\text{1}}$ | $\text{γ}_{\text{1}}$ = 1.8  $\text{γ}_{\text{2}}$ = 1700  $\text{γ}_{\text{3}}$ = 14  $\text{γ}_{\text{4}}$ = 2.3  $\text{γ}_{\text{5}}$ = 14 | $\text{γ}_{\text{j}}\text{ ϵ }\text{[}\text{10}\text{-3}\text{, }\text{10}\text{3}\text{]}$ |
| $\text{ϵ}_{\text{k,}\text{ }\text{j}}$ | The *jth* element of the vector $\text{ϵ}_{\text{k}}$ that denotes the genotype of the *kth* population involved in the consortium (*j* =1 ~ 4; *k* =1 ~ 4). | | N. A. |  |  |  |  |
| $\text{V}_{\text{c}}$ | Volume of the extracellular space | | 10^-3^ L |  | N. A. | N. A. | N. A. |
| $\text{K}_{\text{g}}$ | Half-saturation constant of Monod growth. | | 2×10^-5^ mol⸱L^-1^ | ^26^ | $\text{cp}\text{ }\text{=}\text{ }\text{k}\text{g/}\text{(}\text{Kg}_{\text{i}}\text{k}_{\text{1}}\text{)}$ | $\text{cp}$ = 35 | N. A. |
| $\text{k}\text{g}$ | Maximum consumption rate of P. | | 6.37×10^-4^ mol⸱L^-1^⸱s^-1^ | ^25^ |  |  |  |
| $\text{g}_{\text{k}}$ | Growth rate of the *kth* population (*k* =1 ~ 4). | | N. A. |  | $\text{μ}_{\text{i}}\text{=}\text{g}_{\text{i}}\text{/}\text{k}_{\text{1}}$ | N. A. |  |
| $\text{Y}$ | Yield coefficient for biomass production. | | 10^2^ L⸱mol^-1^ | ^26^ | $\text{y=Y∙}\text{K}_{\text{1}}$ | $\text{y=}$ 9.2×10^-5^ | N. A. |
| $\text{N}_{\text{m}}$ | Carrying capacity of all populations. | | 1.0×10^-3^ L⸱ L^-1^ | ^25^ | $\text{ρ=}\text{N}_{\text{m}}\text{/}\text{V}_{\text{c}}$ | $\text{ρ=}$ 1.0 | N. A. |
| $\text{M}_{\text{k}}$ | Metabolic burdens of the *kth* population (*k* =1 ~ 4). | | N. A. | ^19^ | $\text{m}_{\text{k}}\text{=}\frac{\text{1}}{\text{1}\text{ }\text{+}\left( \sum_{\text{j=1}}^{\text{4}} \text{ϵ}_{\text{k,j}}\text{c}_{\text{j}}\text{e}_{\text{j}} \right)^{\text{h}}}$  $\text{c}_{\text{j}}=\frac{\text{K}_{\text{j}}}{\text{K}_{\text{burd}}}$ (*j* =1 ~ 4) |  | 10^-2^ ~ 1 |
| $\text{K}_{\text{burd}}$ | Coffecient of the metabolic burden. | | Estimated by relative fitness assays (Supplementary Information S1.3.2; Table S1.1) |  |  | Estimated by relative fitness assays (Supplementary Information S1.3.2; Table S1.1) | $\text{c}_{\text{j}}\text{ ϵ (0, 20]}$ |
| *h* | Hill coefficient of the metabolic burden. | | 1.0 |  |  |  |  |
| $\text{X}_{\text{i,0}}$ | Initial biomass of the *ith* population, (*i*=1, 2). | | 1.0×10^-5^ L⸱ L^-1^ | ^26^ | $\text{x}_{\text{i,0}}\text{=}\text{X}_{\text{i,0}}\text{/}\text{V}_{\text{c}}$ | $\text{x}_{\text{i,0}}$ = 0.01 | N. A. |

**Table S2** Definitions of variables and dimensionless methods for our model

| **Variable** | **Definitions** | **Units** | **ND variable** | **ND Value Range** |
| --- | --- | --- | --- | --- |
| $\text{S}_{\text{k,in}}$ | Intracellular concentration of the *substrate* in the *kth* population (*k* =1 ~ 4). | M | $\text{s}_{\text{k,in}}\text{=}\text{S}_{\text{k,in}}\text{/}\text{K}_{\text{1}}$ | 0 ~ 10^5^ |
| $\text{S}_{\text{out}}$ | Extracellular concentration of the *substrate*. | M | $\text{s}_{\text{out}}\text{=}\text{S}_{\text{out}}\text{/}\text{K}_{\text{1}}$ | 0 ~ 10^5^ |
| $\text{I}_{\text{j,}\text{ }\text{k}\text{ }\text{,in}}$ | Intracellular concentration of the *jth* *intermediate* in the *kth* population (*j* = 1 ~ 3; *k* =1 ~ 4). | M | $\text{i}_{\text{i,in}}\text{=}\text{I}_{\text{i,in}}\text{/}\text{K}_{\text{1}}$ | 0 ~ 10^2^ |
| $\text{I}_{\text{j,out}}$ | Extracellular concentration of the *jth* *intermediate* (*j* = 1 ~ 3). | M | $\text{i}_{\text{out}}\text{=}\text{I}_{\text{out}}\text{/}\text{K}_{\text{1}}$ | 0 ~ 10^2^ |
| $\text{P}_{\text{k,in}}$ | Intracellular concentration of the final *product* in the *kth* population (*k* =1 ~ 4). | M | $\text{p}_{\text{i,in}}\text{=}\text{P}_{\text{i,in}}\text{/}\text{K}_{\text{1}}$ | 0 ~ 10^2^ |
| $\text{P}_{\text{out}}$ | Extracellular concentration of the final *product*. | M | $\text{p}_{\text{out}}\text{=}\text{P}_{\text{out}}\text{/}\text{K}_{\text{1}}$ | 0 ~ 10^2^ |
| $\text{X}_{\text{k}}$ | Total biomass of the *kth* population (*k* =1 ~ 4). | L | $\text{x}_{\text{i}}\text{=}\text{X}_{\text{i}}\text{/}\text{V}_{\text{c}}$ | 0 ~ $\rho$ |
| $\text{t}$ | Time | s | $\text{τ=t∙}\text{k}_{\text{1}}$ | 0 ~ 10^8^ |

**Table S3** Strains and plasmids used in this study

| Number | Strains or plasmids | Characterizations | Source |
| --- | --- | --- | --- |
| ***E. coli* strains** | |  |  |
| 1 | *E.coli* Trans5α | F^-^ φ80 *lac ZΔM15 Δ(lacZYA-arg F) U169 endA1 recA1 hsdR17(*r_k_^-^, m_k_^+^*) supE44λ^-^ thi -1 gyrA96 relA1 phoA* | TransGen |
| 2 | *E.coli* HB101 | F^-^ *mcrB mrr hsdS20*(rB mB) *recA13 leuB6 ara-14 proA2 lacY1 galK2 xyl-5 mtl-1 rpsL20*(Smr) *glnV44* | Lab of Professor Lin Min, CAAS |
| ***P. stutzeri* strains** | |  |  |
| 1 | *P. stutzeri* AN0000 | AN1111△*nahC::G*△*nahG::T*△*nahTH::A*△*nahA::G* | ^18^ |
| 2 | *P. stutzeri* AN0001 | AN1111△*nahA::A*△*nahC::C*△*nahG::G* | ^18^ |
| 3 | *P. stutzeri* AN0010 | AN1111△*nahA::T*△*nahC::A*△*nahTH::C* | ^18^ |
| 4 | *P. stutzeri* AN0011 | AN1111△*nahA::T*△*nahC::A* | ^18^ |
| 5 | *P. stutzeri* AN0100 | AN1111△*nahA::C*△*nahG::T*△*nahTH::A* | ^18^ |
| 6 | *P. stutzeri* AN0101 | AN1111△*nahA::A*△*nahG::G* | ^18^ |
| 7 | *P. stutzeri* AN0110 | AN1111△*nahA::A*△*nahTH::G* | ^18^ |
| 8 | *P. stutzeri* AN0111 | AN1111△*nahA::A* | ^18^ |
| 9 | *P. stutzeri* AN1000 | AN1111△*nahC::G*△*nahG::T*△*nahTH::A* | ^18^ |
| 10 | *P. stutzeri* AN1001 | AN1111△*nahC::T*△*nahG::A* | ^18^ |
| 11 | *P. stutzeri* AN1010 | AN1111△*nahC::A*△*nahTH::C* | ^18^ |
| 12 | *P. stutzeri* AN1011 | AN1111△*nahC::A* | ^18^ |
| 13 | *P. stutzeri* AN1100 | AN1111△*nahG::T*△*nahTH::A* | ^18^ |
| 14 | *P. stutzeri* AN1101 | AN1111△*nahG::A* | ^18^ |
| 15 | *P. stutzeri* AN1110 | AN1111△*nahTH::A* | ^18^ |
| 16 | *P. stutzeri* AN1111 | *P. stutzeri* AN10 △*nahW*△*catA*△*Pnah*1*::Ptac*△*Pnah*2*::Ptac*△*nahR:lacI^Q^* | ^18^ |
| 17 | *P. stutzeri* AN10 | Wild type strain | ^1^ |
| **Plasmids** | | | |
| 1 | pRK2013 | Helper plasmid for conjugation; Kan^R^ | Lab of Professor Lin Min, CAAS |
| 2 | pKl8mobsacB | Suicide plasmid vector used for gene knockout; Kan^R^ | Lab of Professor Lin Min, CAAS |

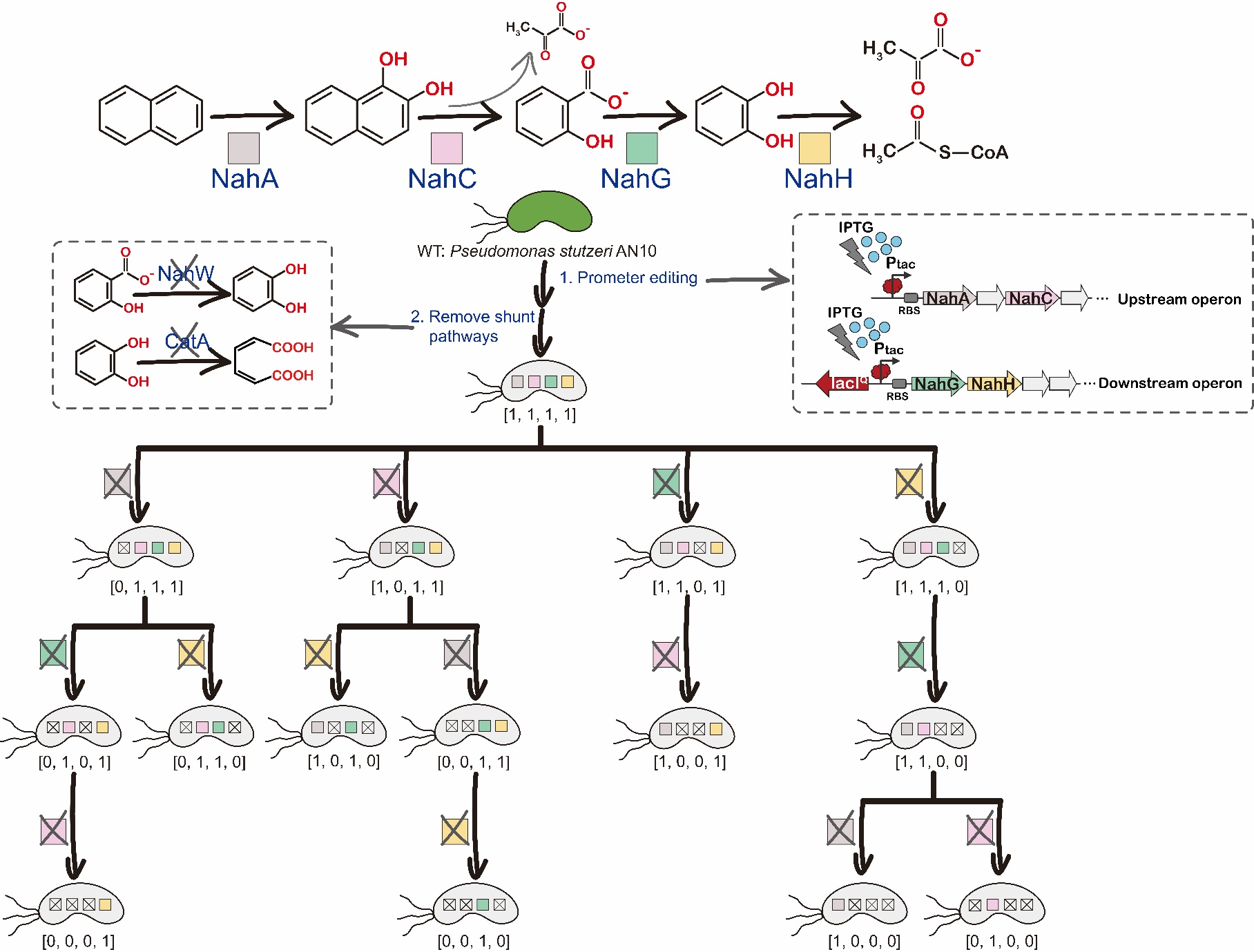

**Figure S1** The workflow of the engineering of the strains used in this study. Related to Figure 1 and Methods. An autonomous strain was engineered by removing the *nahW* gene (encoding a salicylate hydroxylase) and the *catA* gene (encoding a catechol 1, 2-dioxygenase) of *P. stutzeri* AN10. In addition, the original promoters of the two naphthalene degradation operons with an IPTG-induced promoter, *P_tac_*. The derived autonomous strain, namely [1, 1, 1, 1], can degrade naphthalene autonomously, and its genes encoding the enzymes for naphthalene degradation are all located in the two engineered operons. The naphthalene degradation pathway in [1, 1, 1, 1] was further partitioned into four metabolic steps. The four key enzymes responsible for these four steps were encoded by *nahA* gene (encodes a naphthalene dioxygenase), *nahC* gene (encodes a 1, 2-dihydroxynaphthalene dioxygenase), *nahG* gene (encodes a salicylate hydroxylase), and *nahH* gene (encodes a catechol 2, 3-dioxygenase). To construct the mutants that can only perform one or a subset of metabolic steps in naphthalene degradation, the four key genes were knocked out one by one. As a result, forty mutants were further obtained. These strains were named following the same fule as those in Figure S1.

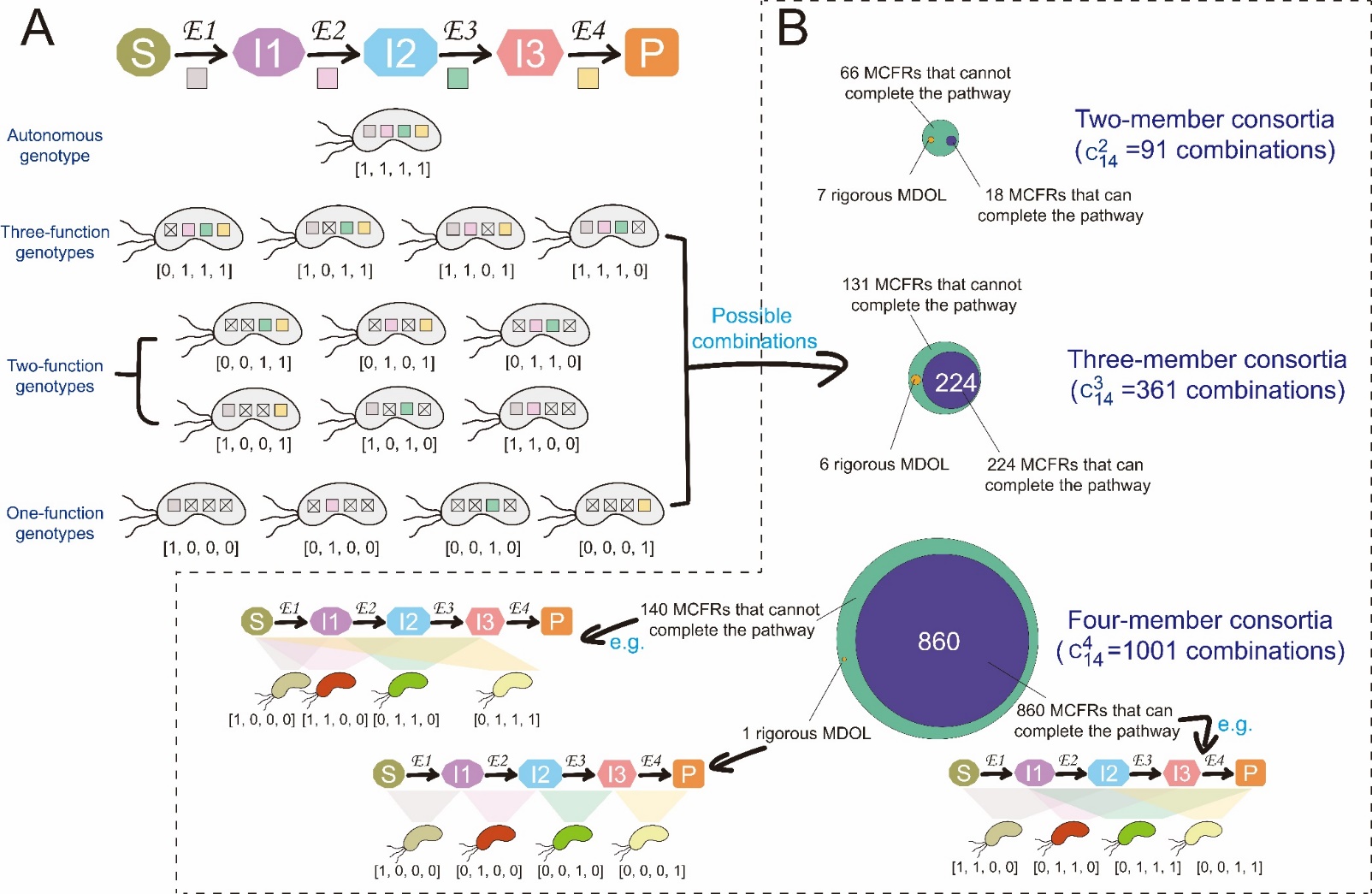

**Figure S2** Schematic diagram of all the genotypes related to performing a four-step pathway (A) and the number of all the possible 2-, 3-, and 4-member consortia composed of these genotypes (B). Related to Figure 1. The genotypes are conceptualized by bit strings containing “0” and “1”. “0” denotes that one genotype lacks the genes responsible for a specific step and “1” denotes that one genotype contains genes encoding the enzymic reactions for a step.

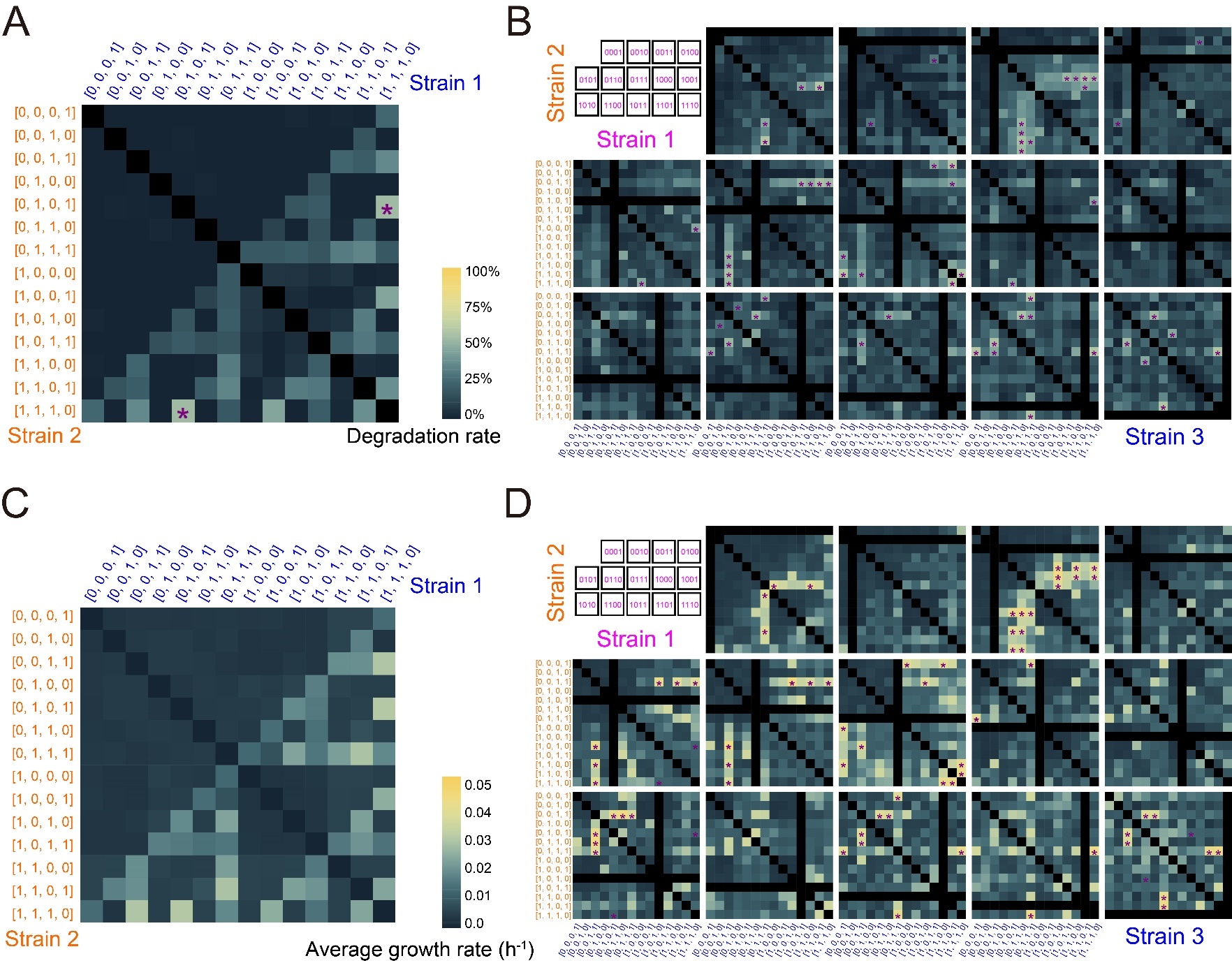

**Figure S3** Degradation rates (A-B) and average growth rates (C-D) of the two-member (A and C) consortia and three-member consortia (B and D). Related to Figure 2. (A) The 14×14 panel shows the naphthalene degradation rate of the two-member consortia composed of the corresponding two strains (blue and orange). (B) The 14×14×14 panels show the naphthalene degradation rate of the three-member consortia. The color intensity in (A) and (B) represents the naphthalene degradation rates after 96-h culture and the scale bar is shown in (A). (C) The 14×14 panel shows the average growth rates of the two-member consortia composed of the corresponding two strains (blue and orange). (D) The 14×14×14 panels show the average growth rates of the three-member consortia. The color intensity in (C) and (D) represents the average growth rates after 96-h culture and the scale bar is shown in (C). In (B) and (D), each 14×14 panel correspond to a fixed strain 1 (violet) against all combination of strains 2 (orange) and 3 (blue). The sequence of strain 1 is indicated in the first panel. All the data were summarized from three independent replicates. The statistical difference between a specific consortium and the autonomous population ([1, 1, 1, 1]) was indicated by a marker “*” involved in the corresponding panel (Double-tailed Student’s T-test, *p* <0.05). In total, one two-member consortium and ten three-member consortia exhibit significantly higher degradation rates than the autonomous population. Twelve three-member consortia exhibit significant growth rates than the autonomous population.

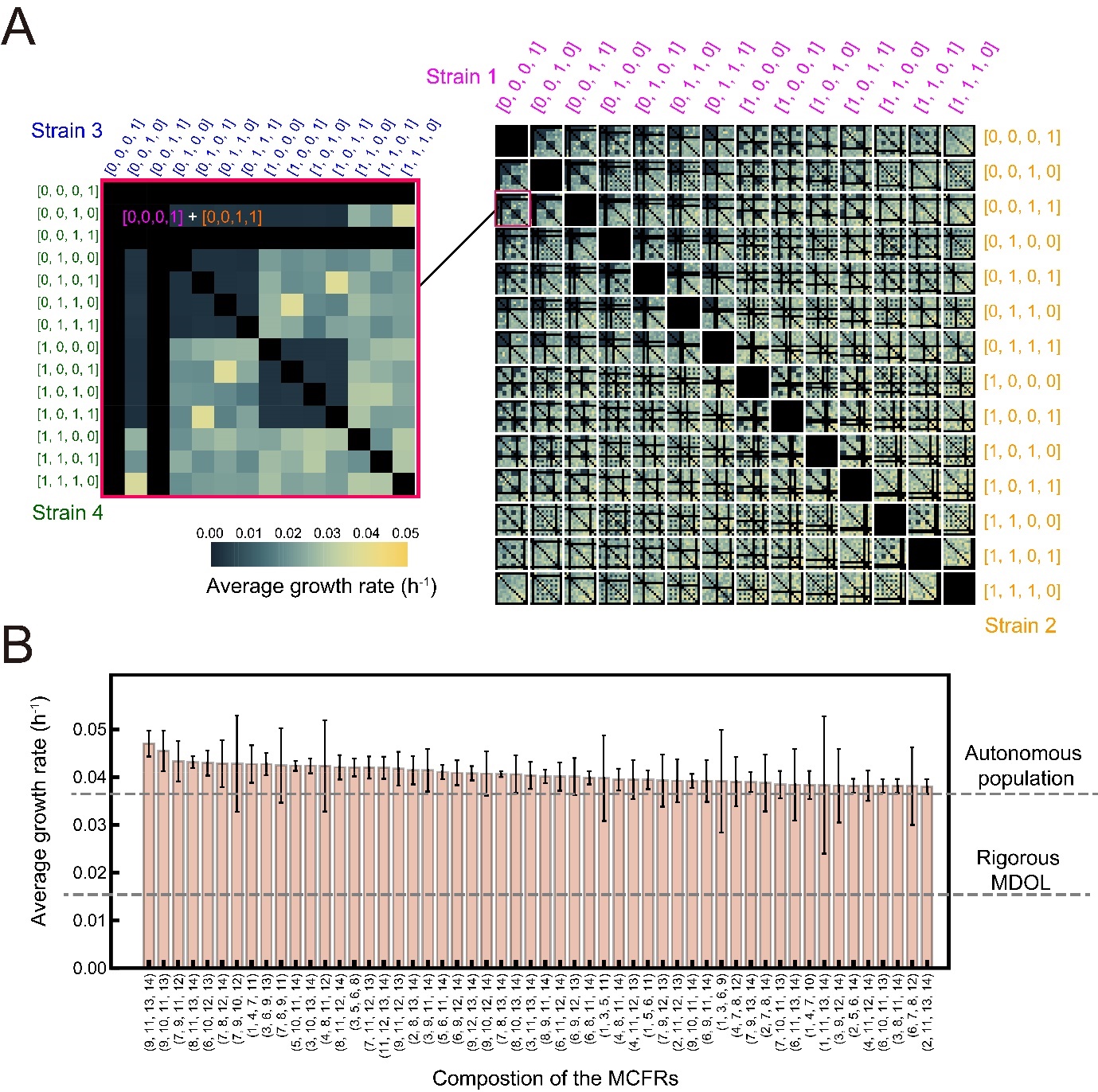

**Figure S4** Average growth rates of the four-member consortia. Related to Figure 2. (A) 14×14×14×14 matrix diagrams show the average growth rates of the four-member consortia. Each 14×14 panel correspond to a fixed strain 1 (violet) and a fixed strain 2 (orange) against all combination of strains 3 (blue) and 4 (green). The color intensity represents the average growth rate across the 144-h culture of the consortia, and the value shown is the average value of the three experimental replicates. (B) Summary of the high-function consortia. Average growth rates of 54 high-function consortia are shown, which are significantly higher than those of the autonomous population and the rigorous metabolic division of labor (MDOL) consortium (Indicated by the grey dashing lines; Double-tailed Student’s T-test, *p* <0.05). The data were summarized from three independent replicates. The consortia are named after the four-number vectors following the same rule as in Figure 2.

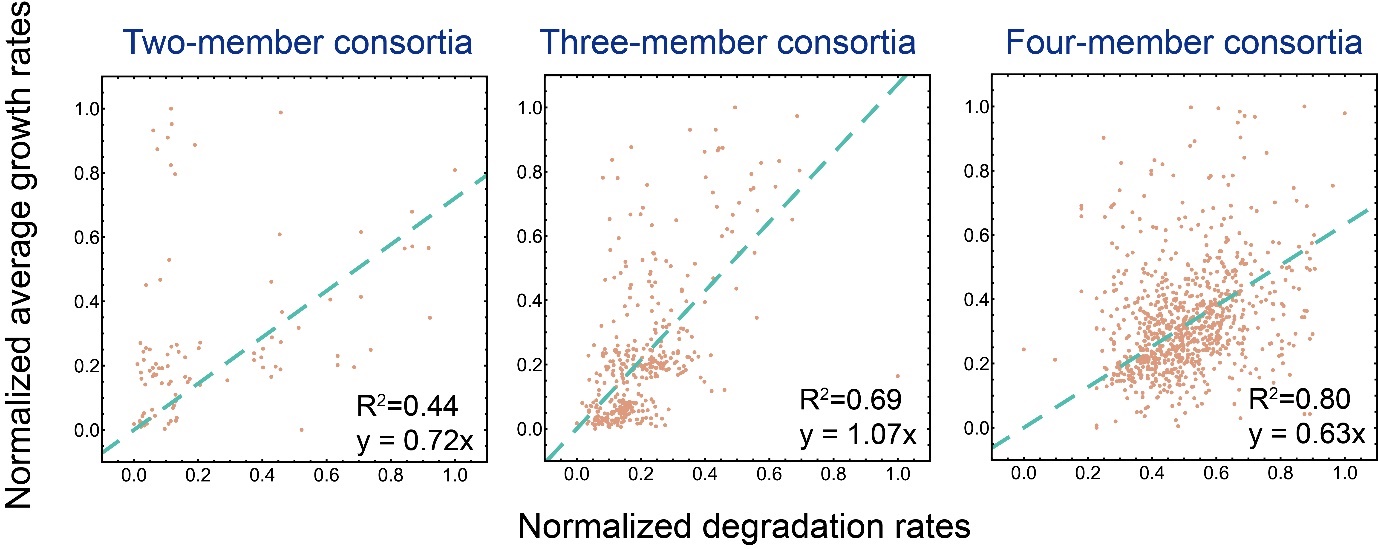

**Figure S5** The correlation between average growth rates and the naphthalene degradation rates of the two-member (*n* = 91), three-member (*n* = 361), and four-member consortia (*n* = 861). Related to Figure 2. The values were normalized according to the maximal and minimal values and the correlations are fitted alongside linear curves (shown by the green dashed lines).

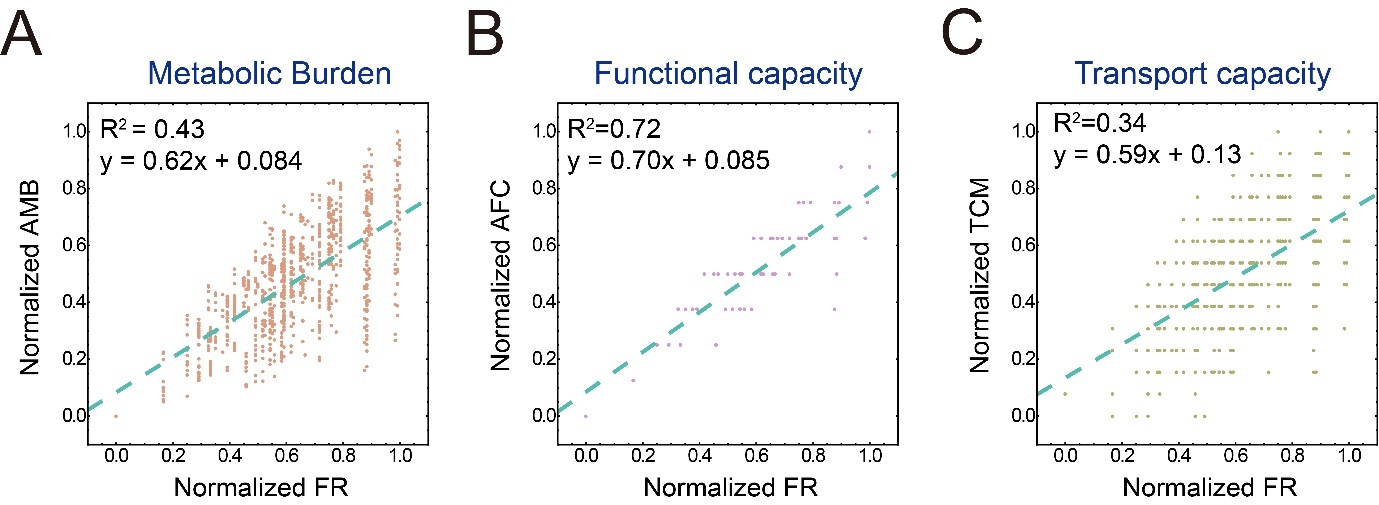

**Figure S6** The correlations between the functional redundancy level (FR) of different consortia and their average metabolic burden of strains (AMB; A), average functional capacity (AFC; B), and transport capacity of metabolites (TCM; C) in our culture experiments of four-member consortia. Related to Figure 3. The correlations are fitted alongside linear curves (shown by the green dashed lines). The definitions of all these indexes are described in Supplementary Information S1.3 and the values were normalized according to the maximal and minimal values.

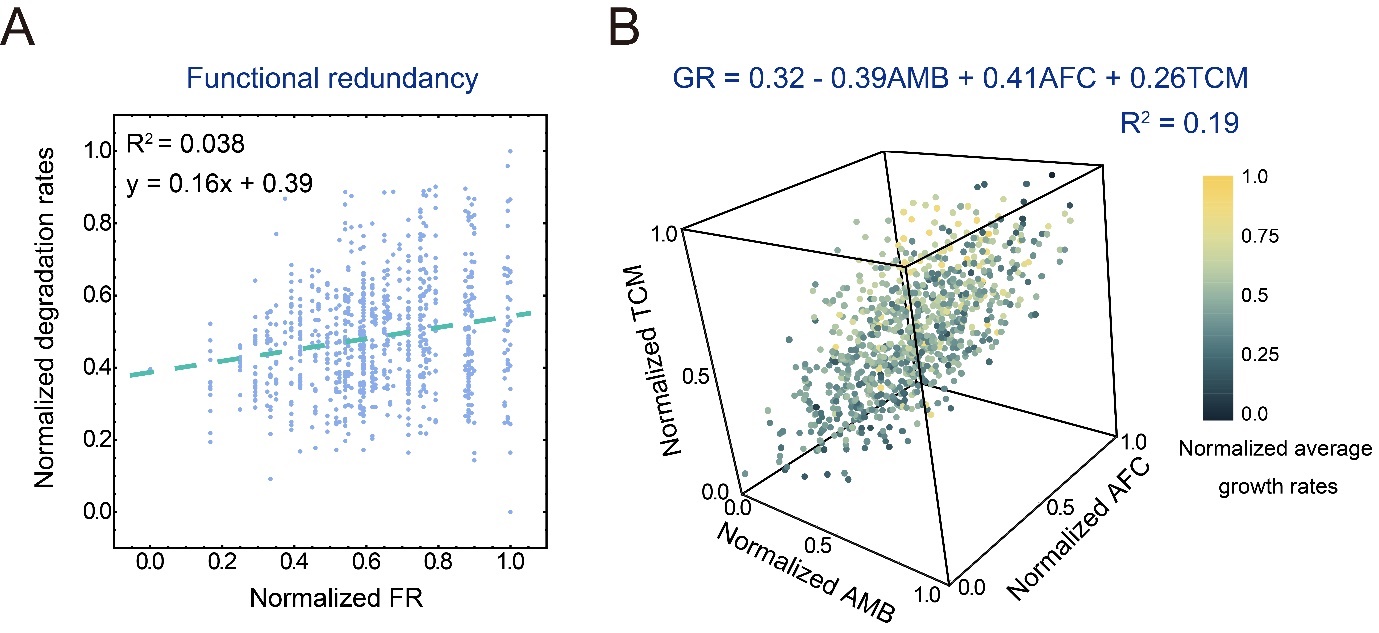

**Figure S7** The growth rates of the engineered consortia can be predicted by the average metabolic burden of strains (AMB), average functional capacity (AFC), and transport capacity of metabolites (TCM). Related to Figure 3. (A) The correlation between the average growth rates of the consortia and their functional redundancy level. The correlation is fitted alongside linear curves (shown by the green dashed lines). (B) The summarized correlations among AMB, AFC, TCM, and average growth rates of the consortia. While the values of AMB, AFC, and TCM are shown by the three axes, the average growth rates are shown by the color intensity. A fitted linear function given at the bottom of the graph indicates the fitted correlation. The definitions of all these indexes are described in Supplementary Information S1.3 and the values were normalized according to the maximal and minimal values.

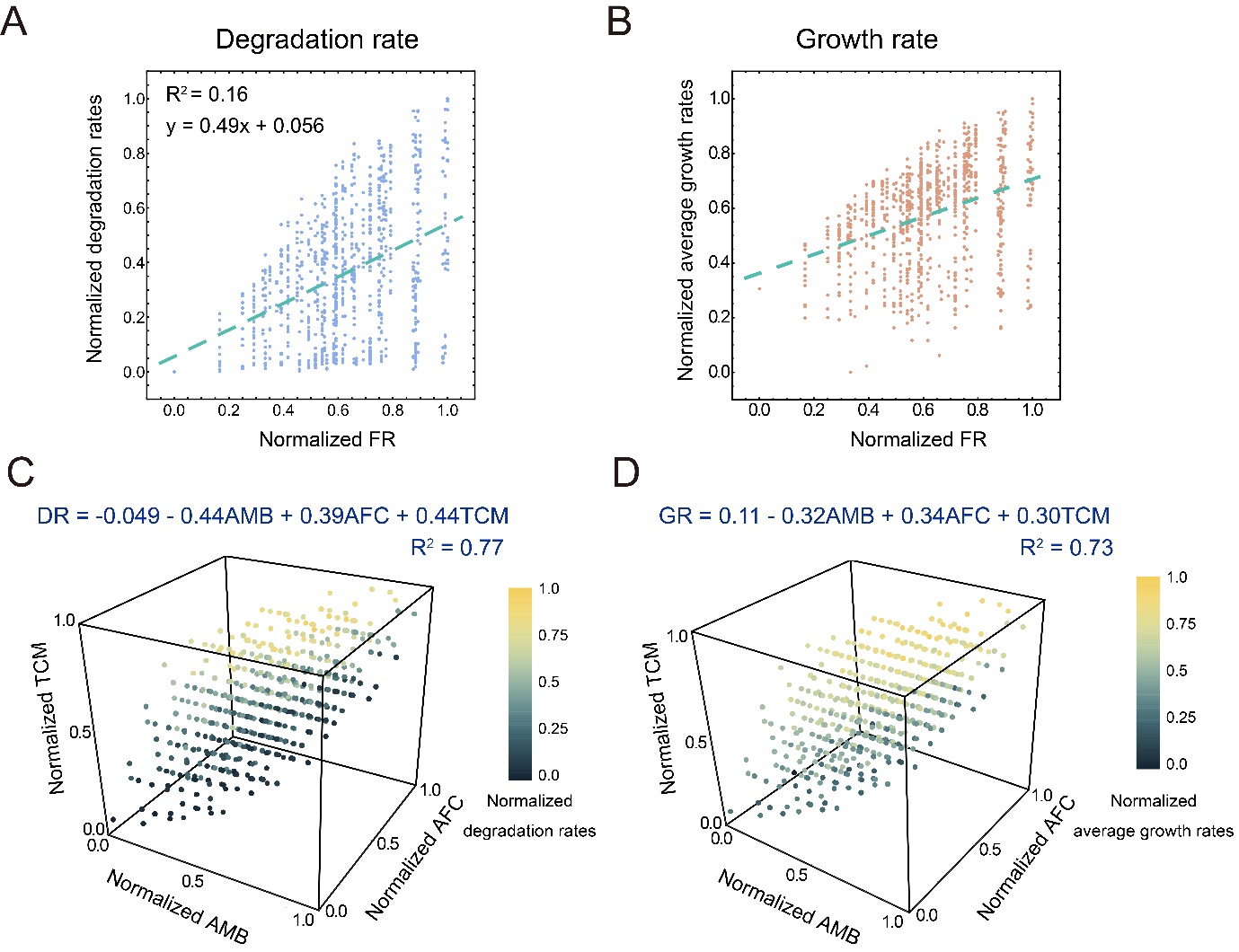

**Figure S8** The degradation rates (A, C) and growth rates (B, D) of the simulated consortia can be predicted by the average metabolic burden of strains (AMB), average functional capacity (AFC), and transport capacity of metabolites (TCM). Related to Figure 4. The simulations were performed using the parameter set derived from our experimental system (Table S1). (A) The correlation between the degradation rates of the simulated consortia and their functional redundancy level. (B) The correlation between the average growth rates of the simulated consortia and their functional redundancy level. The correlation is fitted alongside linear curves (shown by the green dashed lines). (C) The summarized correlations among AMB, AFC, TCM, and the degradation rates of the simulated consortia. (D) The summarized correlations among AMB, AFC, TCM, and the average growth rates of the simulated consortia. While the values of AMB, AFC, and TCM are shown by the three axes, the average growth rates are shown by the color intensity. A fitted linear function given at the left of the graph indicates the fitted correlation. The definitions of all these indexes are described in Supplementary Information S1.3 and the values were normalized according to the maximal and minimal values.

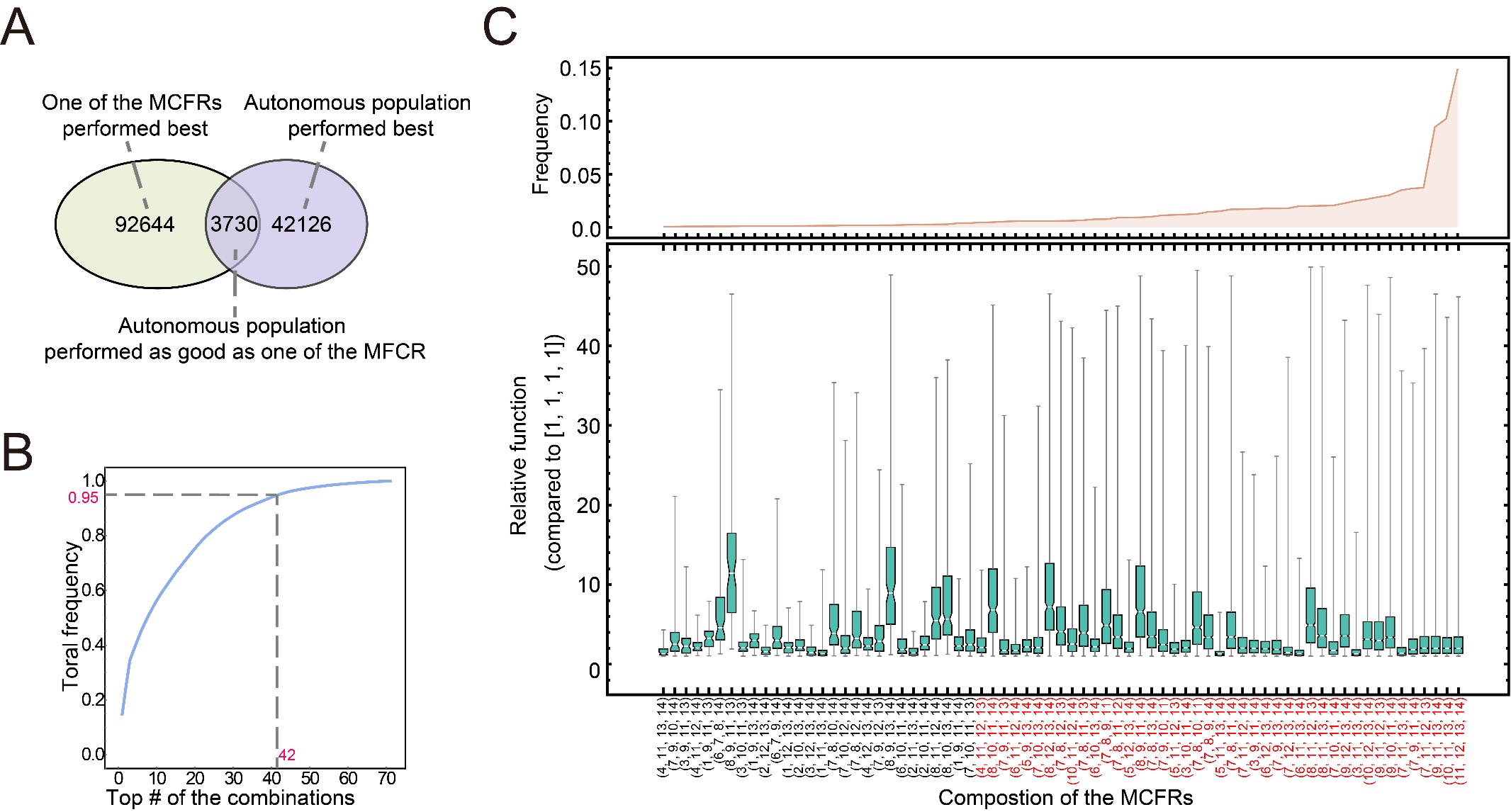

**Figure S9** Simple strategies to screen the optimal synthetic consortia for a given pathway based on the simulated growth rates of different consortia. Related to Figure 5. (A) A Venn diagram indicates the classification of the results of the 138500 simulations into three categories, based on the growth rates of different consortia. (D) Summary of the 71 consortia that possess the highest growth rate in all the simulations. The upper graph shows the frequency of each consortium possessing the highest growth rate in the simulations. The bottom graph shows the relative growth rates of each consortium to that of the autonomous population in all the simulations in which it possesses the highest degradation rate. The consortia are named after the four-number vectors following the same rule as in Figure 2. The names of the consortia with the 42 highest growth rates were labeled red. (E) The accumulated frequency of the consortia with the *N* highest growth rates against the value of *N*. When the consortia with the 42 highest degradation rates are included, their total frequency equals 95 % (indicated by the grey dashed line).

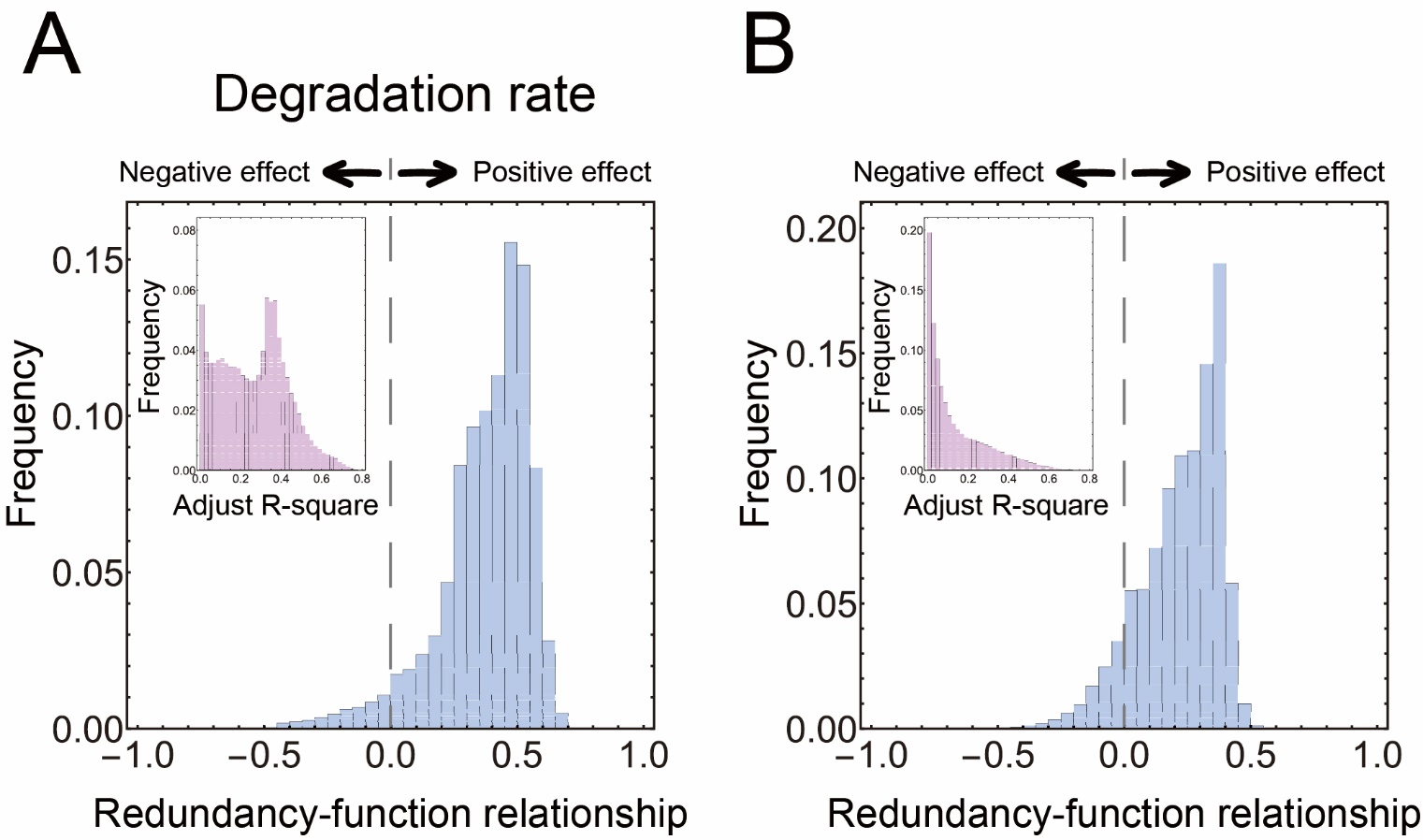

**Figure S10** Distribution of correlation coefficients between the function of the consortia and their functional redundancy level (FR) summarized from the simulations using 135000 parameter sets. Related to Figure 5. (A) the correlation coefficients between the degradation rates of the consortia and their FR. (B) the correlation coefficients between the growth rates of the consortia and their FR. A coefficient value over 0 suggests the functional redundancy level has a positive effect on the degradation rates or growth rates of the consortia, while a coefficient value lower than 0 suggests a negative effect. The inset graphs indicate the distribution of adjusted R-squared values derived from the linear regressions.

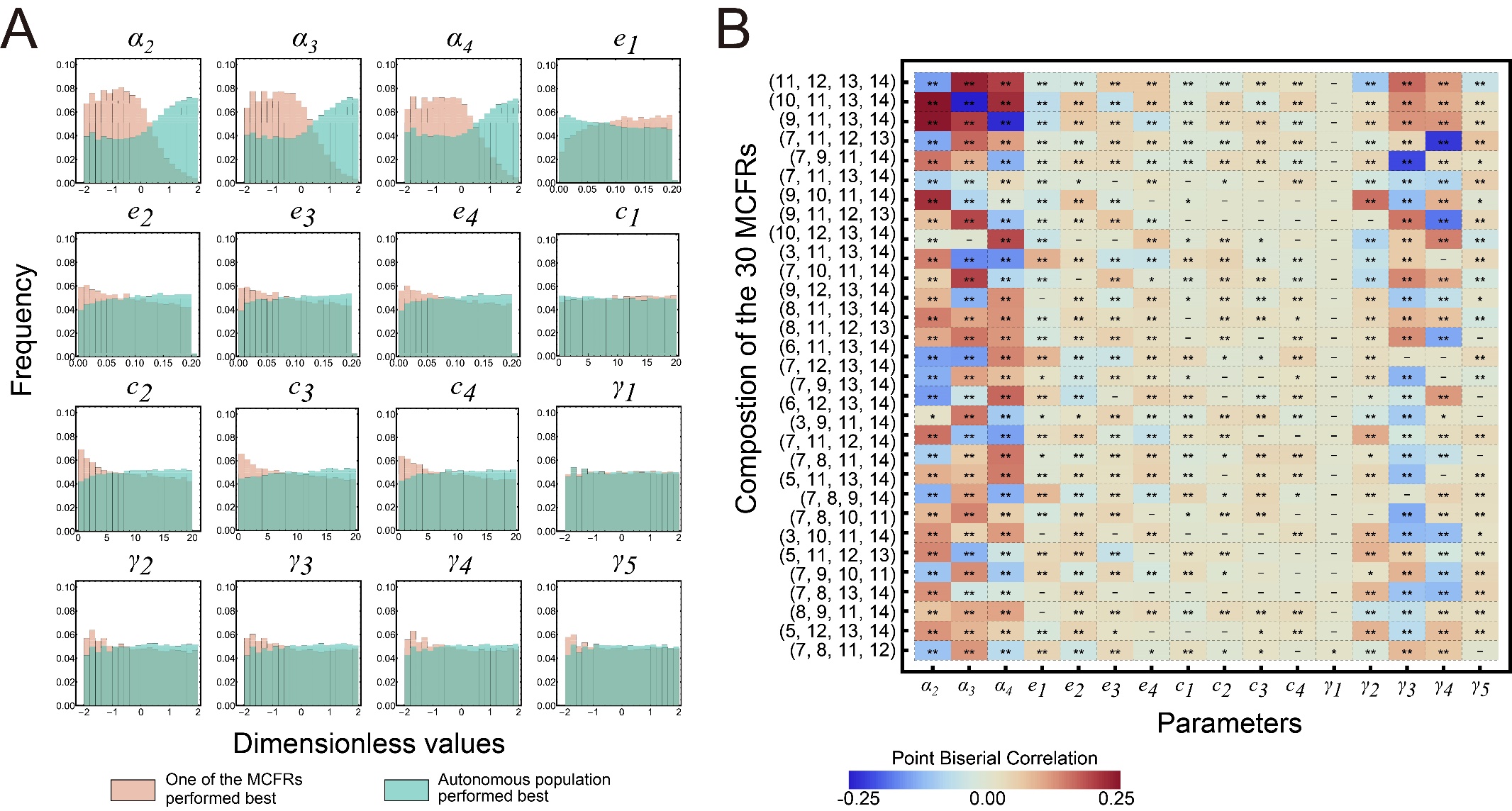

**Figure S11** Key parameters that determine the growth rates of consortia performing metabolic division of labor and possessing functional redundancy (MCFRs). Related to Figure 6. (A) Distributions of two groups of parameter values lead to two different types of simulation results. The light red histograms represent values of the 84248 sets resulting in the simulations in which one of the MCFRs possesses the highest average growth rate. The green histograms represent values of the 52862 sets resulting in the simulations where the autonomous population grows faster than all the consortia. (B) Point Biserial Correlation suggests how the values of different parameters determine whether an MCFR becomes the consortium with the highest growth rate. We set a value of 1 if an MCFR possesses the highest growth rate in a simulation initialized with a parameter set and a value of 0 if it does not grow fastest. As a result, a binary variable is obtained. Then Point Biserial Correlations between the values of each parameter and this binary variable are performed. The values of the derived coefficients are shown by the color intensity. Here, the results of the 30 consortia with the highest frequencies to possess the highest growth rates were shown. The names of these consortia are presented following the same rule as in Figure 2. The markers involved in each grid are derived from Mann-Whitney Tests between the set of the parameters that leads to the corresponding MCFR possessing the highest degradation rate and that leads to MCFR does not perform best. “**”: *p* < 0.0001; “*”: *p* < 0.01; “-”: *p* > 0.01.

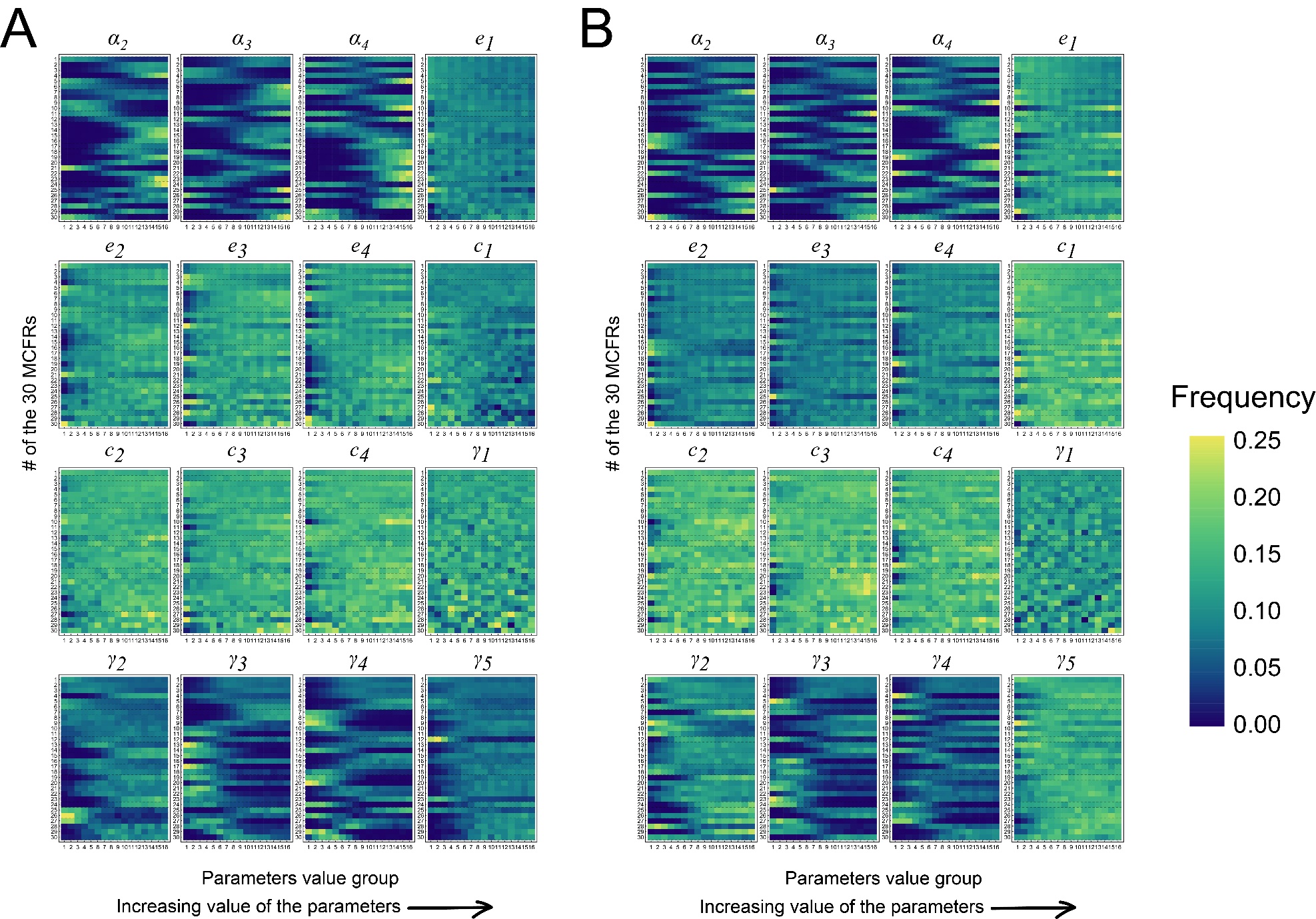

**Figure S12** Distributions of the parameter values in the parameter sets that lead to the simulations in which different consortia possess the highest degradation rate (A) and growth rate (B). Related to Figure 6. We focused on the 30 consortia with the highest frequencies to possess the highest degradation rates (A) and the 30 consortia with the highest frequencies to possess the highest growth rates (B). For a given consortium, we selected the simulations in which this consortium possesses the highest degradation rates or growth rates and analyzed the distributions of the parameter values of these simulations. The parameter values were classified into 16 groups and the frequencies of the simulations involved in each group are shown by the color intensity.

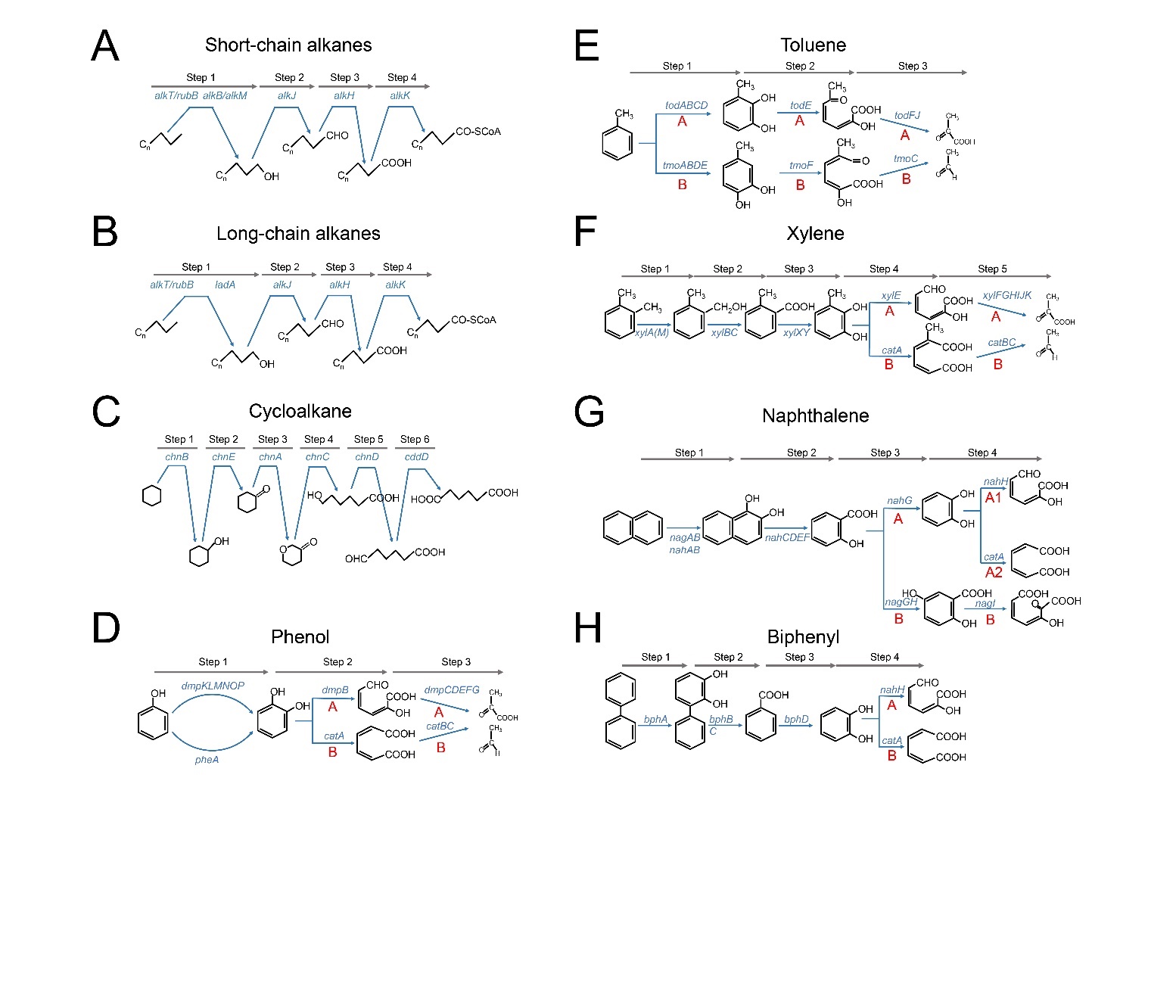

**Figure S12** Deconstruction of the degradation pathways of eight hydrocarbons into three to six steps. Related to Figure 7.

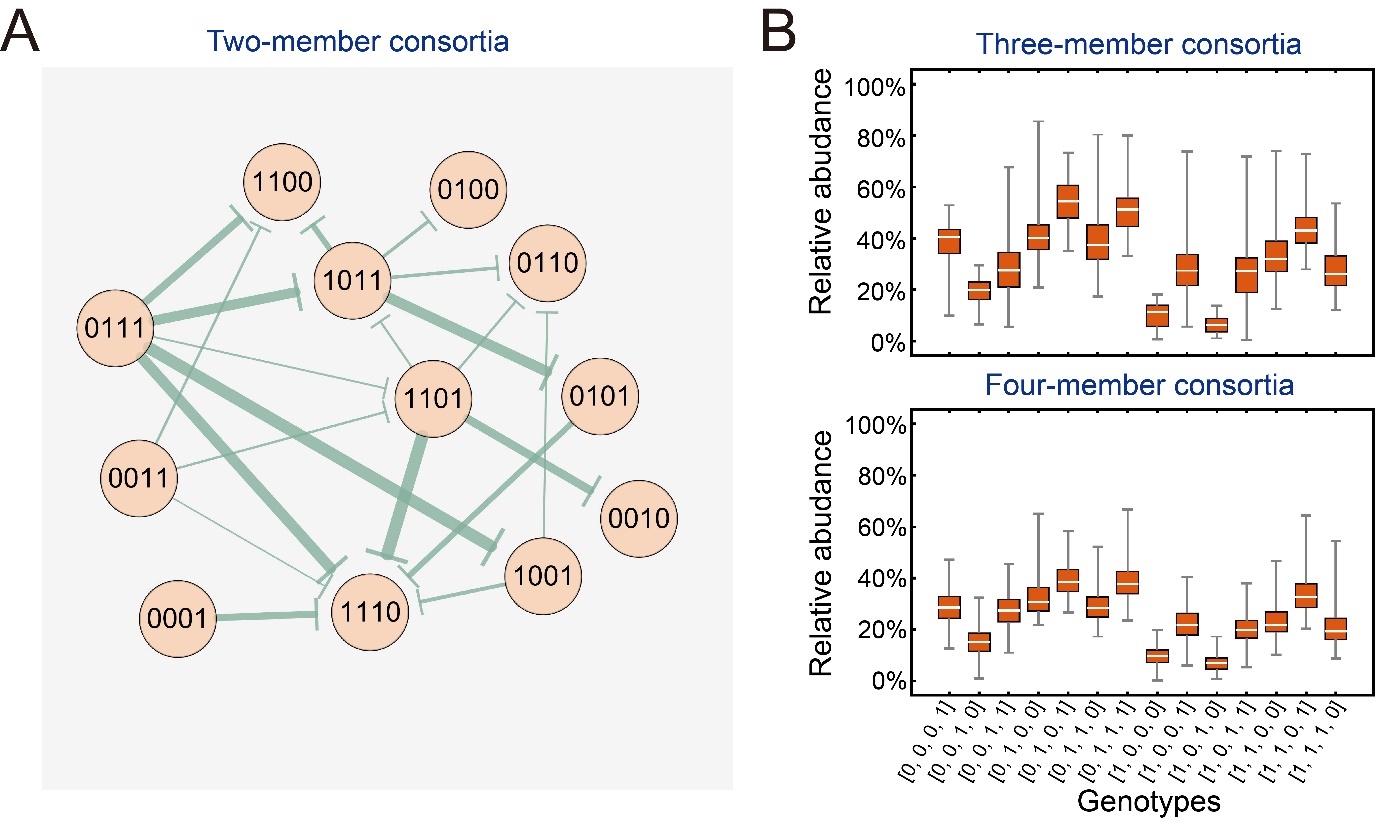

**Figure S14** Relative abundances of different genotypes in the culture experiments of different consortia. Related to Figure 2. (A) Analysis of pairwise correlation of different strains based on the relative abundance of the two strains in their co-culture (e.g., the culture of the two-member consortia). The starting side of the arrow is the strain with the higher abundance, while the ending side denotes the strain with the lower abundance. The thickness of the arrow represents the ratio of their abundance. (B) Distribution of the relative abundance of different genotypes in the culture of all three-member consortia (the upper graph) and four-member consortia (the below graph). The relative frequencies were measured after 96-h culture of the consortia, following the protocols described in Supplementary Information S1.2. All the data were summarized from three independent replicates.
